# The Peptonizer2000: bringing confidence to metaproteomics

**DOI:** 10.1101/2024.05.20.594958

**Authors:** Tanja Holstein, Pieter Verschaffelt, Tim Van Den Bossche, Simon Van de Vyver, Lennart Martens, Bart Mesuere, Thilo Muth

## Abstract

Metaproteomics, the large-scale study of proteins from microbial communities, faces challenges in identifying species due to similarities in protein sequences across different organisms. Current methods often rely on simple counting of matches between proteins and taxa, which can lead to low accuracy. We introduce the Peptonizer2000, a new tool that uses advanced modeling to provide more precise taxonomic identifications along with confidence scores. It combines peptide scores from any proteomic search engine with peptide-to-taxon links from the Unipept database. By applying statistical models, the Peptonizer2000 improves taxonomic resolution and delivers more reliable results. We validate its performance using publicly available datasets, demonstrating its ability to produce high-confidence identifications. Our results suggest that the Peptonizer2000 improves the specificity and confidence of taxonomic assignments in metaproteomics, providing a valuable resource for the study of complex microbial communities.

## Introduction

Microbial communities, also known as microbiomes, can be found in diverse environments such as ocean water,^1^ biogas plants,^2^ and the human gut.^3^ Recent technological and bioinformatic advances have enabled new discoveries such as the ability of microbiomes to use carbon monoxide in certain marine worms,^4^ insights into the composition of certain environmental microbiomes,^5^ the alteration of the gut microbiome depending on diet,^6^ and, more generally, the role microbiomes play in health and disease of humans^7^ and animals.^8^ Metaproteomics is a growing field that studies microbiomes by directly analyzing their proteins to investigate microbiome function and taxonomic composition.^9–11^ Tandem mass spectrometry (MS/MS), a technology from traditional proteomics, is employed to analyze proteins in microbiome samples. However, microbiome samples are more complex to handle and understand than single organism samples, which poses challenges in experimental and bioinformatic setups.^12,13^ One key objective of metaproteomics is to identify the microorganisms present in the sample along with their roles. This task is complicated by the fact that proteins often share similarities in their sequences and that they can be linked to multiple groups of organisms.^14^ In very complex samples, identifying specific species or even families of microorganisms can be challenging.

Several bioinformatic tools have been developed to identify taxa and functions in microbiome samples. Among the most popular software tools are MetaProteomeAnalyzer,^15^ iMetaLab,^16^ the Galaxy framework,^17^ and Unipept.^18^ These workflows often rely on heuristics, such as peptide-taxon match counting, to determine the present taxa. Commonly, taxa are considered present if at least two, or sometimes three, unique peptides mapping to the taxon are detected.^19–21^ These methods disregard underlying uncertainties inherently linked to any data collection method and may suggest a false sense of confidence in the obtained results. Additionally, when a peptide cannot be assigned to a specific species, it is mapped to the lowest taxonomic level to which it is specific, reducing the taxonomic resolution of the analysis. This taxonomic level is referred to as the lowest common ancestor (LCA).^22^ The LCA approach is key to the taxonomic identification performed by Unipept. Over recent years, the number of taxa and associated proteins in publicly available databases like UniProt has steadily increased from 51 million in 2014 to 245 million in 2024.^23^ This growth has been shown to decrease the specificity of peptides at lower taxonomic levels, thus reducing the taxonomic resolution of the LCA approach.^18^

Addressing these issues requires more sophisticated taxonomic identification algorithms. Mi-CiD^24^ is one such method, providing integrated metaproteome analysis supported by rigorous statistical methods for accurate taxonomic identification. However, MiCiD uses its own built-in database search for mass spectrum identification, which is tightly linked to its taxonomic assignment process. As a result, it is not compatible with outputs from other widely used search engines like X!Tandem^25^ or rescoring tools like MS2Rescore^26^ and is restricted to data-dependent acquisition (DDA) spectra only.

In contrast, Unipept can flexibly process peptide lists from any search engine but does not incorporate peptide scores in its analysis. Instead, it relies on counting peptides assigned to the LCA for taxonomic inference, which limits its precision.

None of the aforementioned software solutions combine advanced statistics, straightforward usability, comprehensive visualization options, and compatibility with any preferred search engine. Furthermore, they lack support for both DDA and data-independent acquisition (DIA) data despite DIA’s increasing popularity in recent metaproteomics studies.^19,27,28^

## Our contribution

We introduce the Peptonizer2000, a novel workflow for taxonomic identification of metaproteome samples. Unlike previous approaches, the Peptonizer2000 models the joint probability distribution of all detected peptides and the potentially present taxa using a graph representation. As an alternative to arbitrary heuristics, it uses a Bayesian approach to compute the presence of taxa by calculating their marginal probabilities. This results in taxonomic identifications with associated confidence scores. We previously developed PepGM,^29^ a graphical model-based workflow for the taxonomic identification of single-virus proteome samples. In that work, we addressed the challenge of strong protein sequence homology between viral strains, which can make accurate strain-level identification difficult, an issue that is analogous to the one faced here when identifying the species present in microbiomes. In this study, we have shown that using a more advanced statistical model leads to more accurate taxonomic assignments. Building on this, the Peptonizer2000 integrates statistical modeling for taxonomic assignments, incorporating peptide scores from proteomic search engines along with peptide-taxon mapping from Unipept. This allows the Peptonizer2000 to provide high-resolution taxonomic identification for microbiome samples, including probability scores for each taxon. By doing so, the Peptonizer2000 adds confidence to taxonomic assignments that were, until now, reliant on discretionary PSM cutoffs.

We show that our new approach can compute taxonomic confidence scores at user-selected taxonomic levels for various metaproteomics samples using different background databases. Comparing Peptonizer2000’s probabilistic identifications with Unipept’s PSM-based counts, we demonstrate how these confidence scores offer valuable evidence to distinguish between truly present taxa and spurious taxon-peptide matches.

## Results

### Overview of the Peptonizer2000 workflow

A broad overview of the Peptonizer2000 workflow is shown in Figure 1, which consists of three main steps: first, querying all candidate peptides in the Unipept database and collecting candidate taxa based on weighted PSMs; second, constructing the graphical model, performing belief propagation, and conducting a grid search through graphical model parameters; and third, evaluating the parameters, followed by output and visualization of the results. These steps are adapted from our previously developed taxonomic identification tool, PepGM.

**Figure 1:**
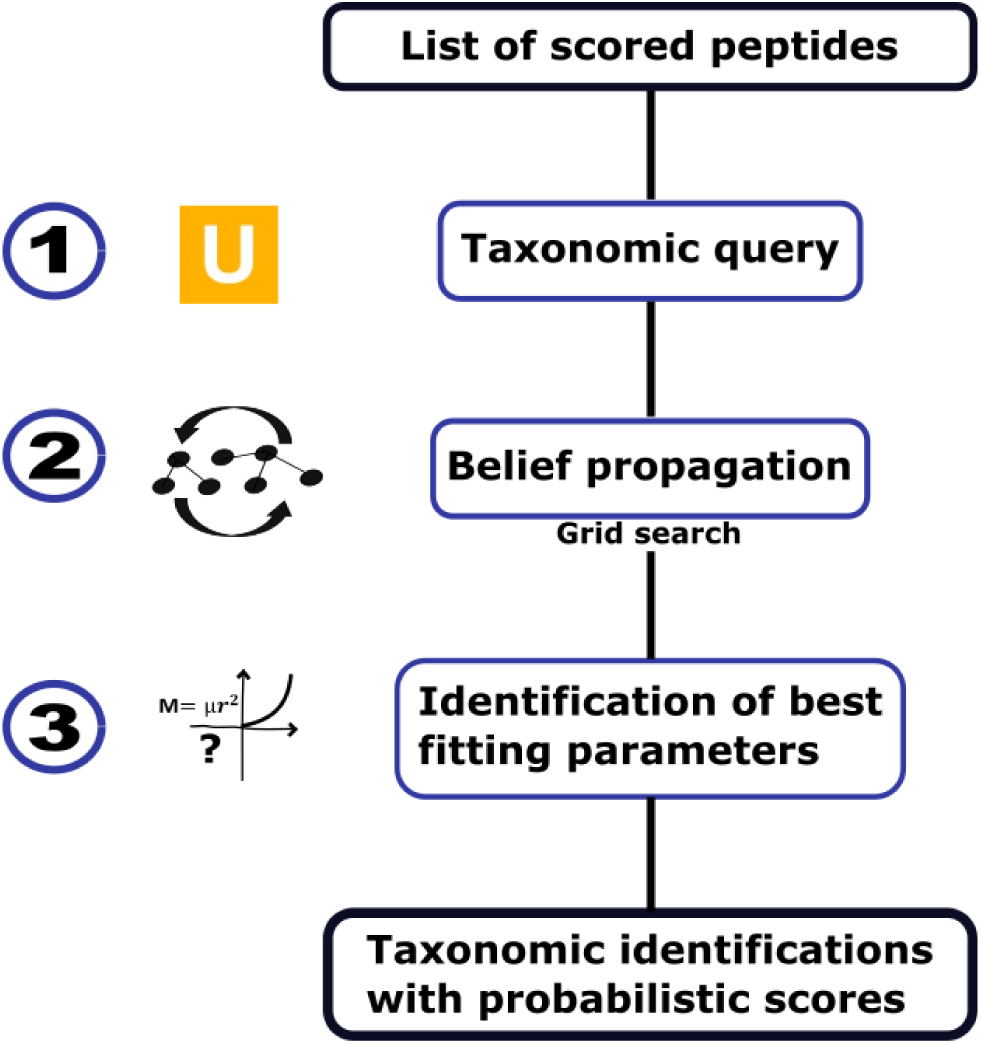
Overview of the Peptonizer2000 workflow. A list of scored peptides has to be provided as input. (1) These peptides are queried for taxonomic information using Unipept. (2) Using a factor graph, the probabilities at the peptide level are propagated to the taxa. Model parameters are evaluated through a grid search. (3) The optimal parameters for each sample are determined using a custom metric, resulting in taxonomic identifications with associated confidence scores.

Snakemake^30^ is used as the workflow management system, while all scripts are written in Python and developed and tested for the Linux operating system and Python version 3.10. All additional Python packages used are listed in the supplementary materials.

The Peptonizer2000 source code is open source and is available under https://github.com/BAMeScience/Peptonizer2000. The version of the Peptonizer described in this work is available under the ’peptonizer2000 manuscript’ tag.

We provide a detailed description of the workflow and algorithms used in the Methods section.

### Taxonomic identification results for samples with known composition and varying complexity

To highlight the benefits of probabilistic scoring for taxonomic identification and to demonstrate the accuracy of the Peptonizer2000, we analyzed a selection of publicly available datasets using both Unipept and the Peptonizer2000.

#### Taxonomic composition of lab-assembled mixtures

For initial testing, we used lab-assembled mixtures with a defined taxonomic composition, providing a clear ground truth for the species present in the samples. This means that we can clearly assess the Peptonizer2000’s performance: if all taxa known to be in the ground truth are correctly reported, and only those, the Peptonizer2000 performs well. We selected two publicly available datasets: one (referred to as the SIHUMIx sample) from the CAMPI study,^31^ a large, cross-laboratory metaproteomic study from the Metaproteomics Initiative,^32^ and the other from a study that assesses methods for estimating biomass contributions in microbiomes.^33^ We began our analysis with the provided spectral and database files from the respective datasets, performing the database search and rescoring as detailed in the Materials section.

#### Taxonomic composition results for the SIHUMIx sample

We analyzed the taxonomic composition of the samples S03, S05, S07, S08, and S11 from the SIHUMIx dataset of the CAMPI study. SIHUMIx is composed of *Bacteroides thetaio-taomicron*, *Anaerostipes caccae*, *Escherischia coli*, *Lactoplantibacillus plantarum*, *Clostridium butyricum*, *Thomasclavelia ramosa*, *Blautia producta* and *Bifidobacterium longum*. As a reference database, we used a sample-specific SIHUMIx database, provided alongside the MS/MS data in the original study. This database includes *Blautia pseudococcoides* as the representative species of the genus *Blautia*, as this is the available reference proteome. In the following, we therefore also accept *Blautia pseudococcoides* as a valid taxonomic identification result.

Figure 2A shows the analysis results for the top-scoring taxa in sample S07. The Peptonizer2000 correctly identifies all present taxa as top-scoring, with probability scores ranging from *p* = 1 to *p* = 0.9. Additionally, all present taxa are accurately represented among those with the highest number of unique peptides. The results also demonstrate several notable attributes of both Peptonizer2000 and Unipept, which we will explain in the following. The Unipept results show a very high range in the number of detected peptides (between 2 and 1399) for the taxa present in the sample. From the Unipept results alone, selecting the present taxa with arbitrary criteria like ’a minimum of 2 unique peptides per taxon’ is hard to justify, even though in the present case, it would lead to correct identifications. This type of cutoff is very common in metaproteomic data analysis.^19–21,34,35^

**Figure 2:**
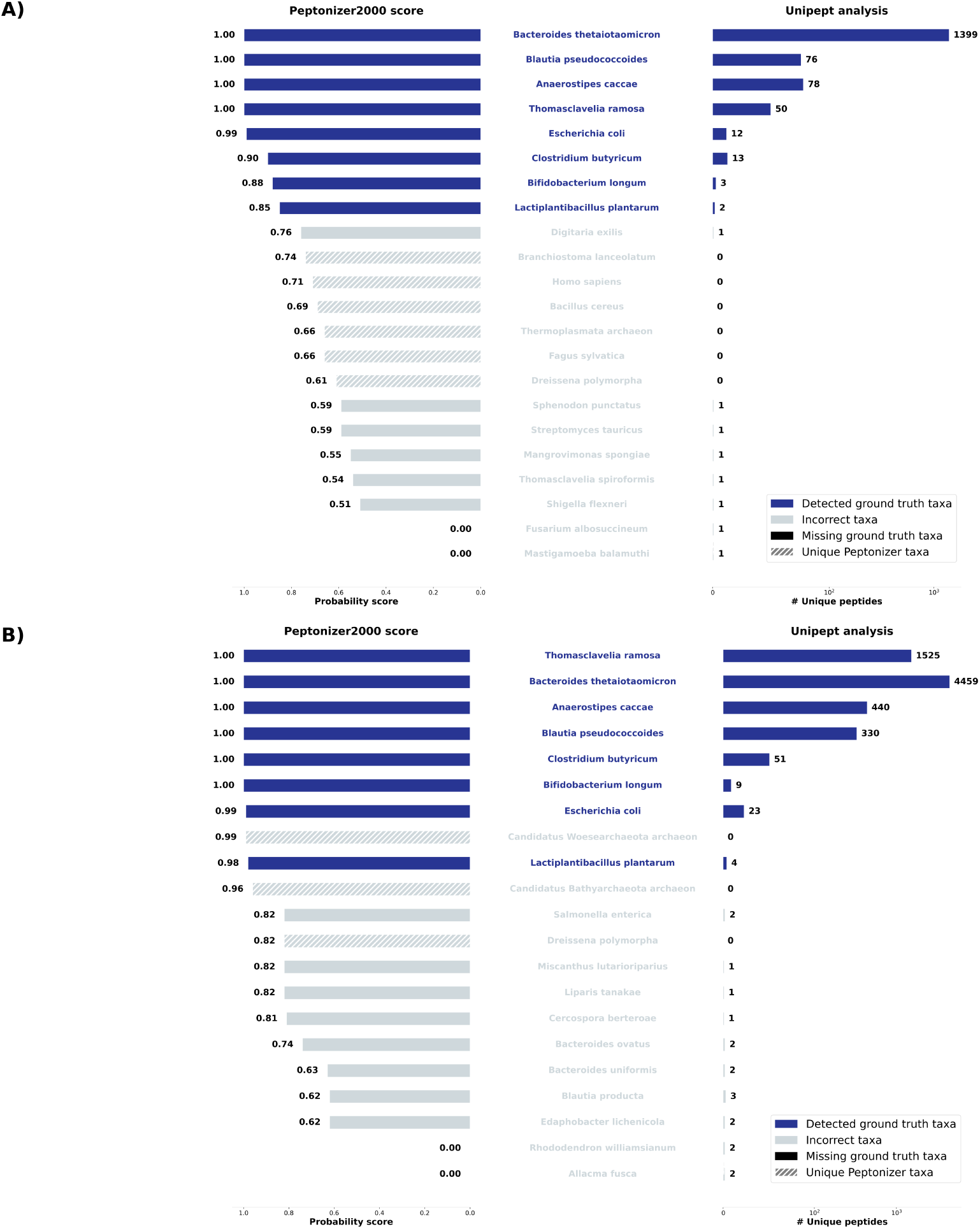
Taxonomic identification results for the SIHUMIx samples (A) S07 and (B) S11. The left panels display the results from Peptonizer2000, including corresponding probability scores, while the right panels show results based on unique peptides from Unipept. Dark blue shades indicate correct taxonomic identifications. The two software tools demonstrate a high degree of overlap in their results.

The Peptonizer2000 attributes all the highest scores to the correct taxa. Since they represent a probability distribution, introducing a ’cut-off’ probability for present taxa does not make sense mathematically but will often be necessary for practical reasons. Here, looking at the results from Unipept and Peptonizer2000 conjointly, a posterior taxon probability of about *p* = 0.9 could be suggested. Additionally, combining the results from both tools yields the most accurate outcomes: the eight taxa with the highest number of unique peptides also received high Peptonizer2000 scores, allowing for confident identification of their presence. None of the taxa with a single unique peptide appear among the higher-scoring taxa in the Peptonizer2000 results, indicating their absence with a high degree of confidence. The next higher scoring taxon in the Peptonizer2000 results is *Branchiostoma lanceolatum*. In the Peptonizer2000 graph, this species has 51 shared peptides attributed to it, which is why it still obtains a relatively high probability score of ≈ 0.85. However, looking at the Unipept results, one can see that it has no unique peptides. With the Unipept and Peptonizer2000 scores together, it can, therefore, safely be identified as absent.

Figure 2B displays the results for the SIHUMIx sample s11. All taxa known to be present in the lab assembled mixture are detected as present by the Peptonizer2000 with high confidence. However, the Peptonizer2000 incorrectly identifies *Candidatus Woesearcheota archaeon* and *Candidatus Bathyarcheota archaeon* as present. These taxa are represented by one unique peptide and share many peptides with taxa that are actually present. Most importantly, the *Candidatus* in the taxon name refers to putative, yet uncultured eukaryotic taxa, and the records associated to *Candidatus* taxa are known to be error-prone and disorganized.^36^ Researchers can thus easily identify it as an erroneous classification. In the future, *Candidatus* could be included into the prefiltering that is already being performed on the Unipept side, hence the absence of any *Candidatus* taxa in the Unipept results. The same observations for sample S07 apply to the remaining high-scoring taxa identified by Peptonizer2000 and the unique peptides detected by Unipept in the current sample. Here, Peptonizer2000’s probability scores support using a threshold of three unique peptides per taxon for confident identification. Additionally, a posterior probability of *p* = 0.9 again appears to be a reliable indicator of presence. Other taxa in the Unipept results, including those with two unique peptides, show lower probability scores, suggesting they are less likely to be present.

Similar conclusions can be drawn from the Peptonizer2000 and Unipept results for the samples S03, S05, and S08 (Supplementary Figures 1-3).

#### Taxonomic composition results for more complex samples

To assess the performance of the Peptonizer2000 on complex samples, we analyzed a dataset from a previously studied lab-assembled mixture containing up to 32 microorganisms, as reported in a prior metaproteomic study.^33^ We examined samples U1, C1, and P1, which were constructed with uneven taxonomic abundance, protein, and cell number (U1), equal cell amount (C1), and equal protein amount (P1).

Figure 3A shows the Peptonizer2000 identification results for sample U1 in a treeview diagram, displaying all taxa with a probability score above 0.8. Additionally, Figure 3B illustrates the Peptonizer2000 attributed scores for all taxa scoring above 0.8 together with the 30 taxa with the highest number of unique peptides using Unipept.

**Figure 3:**
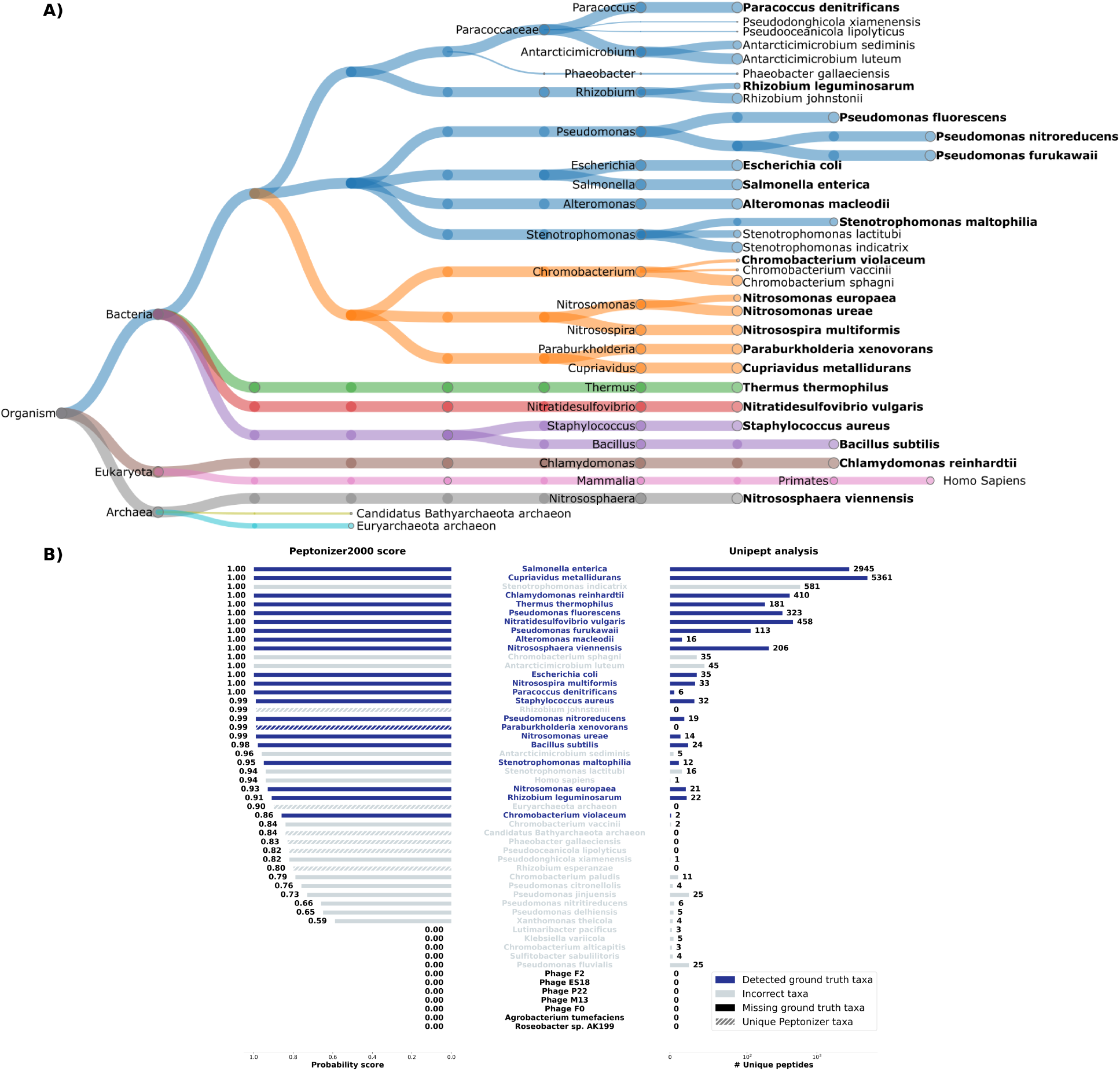
(A) Taxonomic tree view of the Peptonizer2000 results for lab-assembled sample of 28 different species, filtered with a probability score above 0.8. Node sizes are proportional to probabilities attributed by the Peptonizer2000. Correct taxonomic identifications are displayed in bold font. (B) Comparison of the Peptonizer2000 probability scores (left) and the number of unique peptides detected per taxon using Unipept (right). Correct identifications are shaded in darker blue.

Of the 32 potentially present taxa, 28 were included in the lab-assembled mixture by the original study designers. Among these, 5 are phages, which we did not detect, likely due to their very low protein amounts and the filtering steps currently applied by Unipept. Additionally, taxonomic annotations for viruses and phages in reference databases are often less comprehensive and less curated than those for bacteria and eukaryotes.^37^ Of the remaining 23 microorganisms, 21 were correctly identified. The two missing taxa are *Roseobacter Sp. 199* and *Agrobacterium tumefaciens*. The species *Roseobacter Sp. 199* is currently absent in the NCBI taxonomy, which could explain its absence from the results. Some erroneous taxa received scores above 0.8. However, most of these are closely related to the present species. For example, *Rhizobium johnstonii* was incorrectly identified along with the correct *Rhizobium leguminosarum* from the same genus. Similarly, the two *Pseudomonas* species, *indicatrix* and *lactitubi*, were wrongly identified. The Peptonizer2000 cannot distinguish between species within these genera based on the detected peptides.

The comparison between the Peptonizer2000 and Unipept results in Figure 3 highlights the advantage of integrating their outputs. The Peptonizer2000 scores provide valuable insights into the presence or absence of taxa with unique peptides, especially considering the broad range of unique peptides mapped to identified taxa (from 6 to 5000). In the current sample, the Peptonizer2000 scores confirm the presence of *Paracoccus denitrificans* and the absence of *Pseudomonas nitrireducens*, even though both taxa have six unique peptides attributed to them. However, some false positives persist. For example, *Antarctimicrobium sediminis* obtains a high probability score of 0.96 and has five unique peptides despite not being present in the sample. Additionally, two archaeal species are identified as false positives in the Peptonizer2000 results, one of which includes *Candidatus* in its name, suggesting potential reference database issues. Since archaea are frequently misidentified, implementing an optional filter for ’archaea’ taxa could be a useful feature in future updates.

The taxonomic analysis of samples C1 and P1 is presented in the supplementary materials (Figures 4 and 5). While the number of correct identifications is lower—18 for community C1 and 16 for community P1—the findings regarding the complementary value of unique PSMs from Unipept and Peptonizer2000 scores still hold true. Notably, for both C1 and P1, the species Agrobaterium fabrum (heterotypic synonym of *Agrobacterium tumefaciens*) is correctly detected with a score of 1 even though only one or no unique peptide was detected by Unipept.

**Figure 4:**
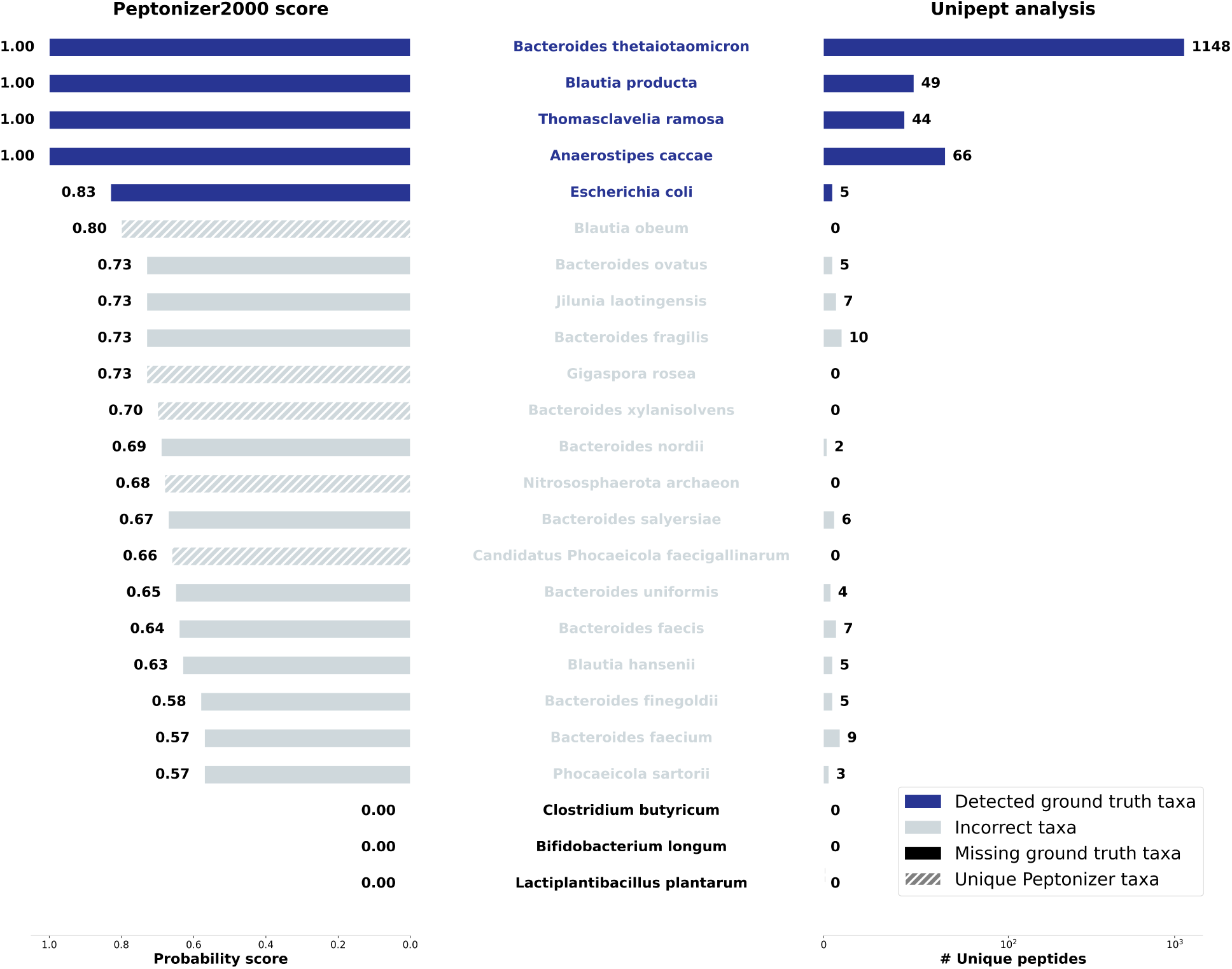
Taxonomic identification of the SIHUMIx sample s07 searched against the IGC reference catalog. The left panel shows results using Peptonizer2000 with corresponding probability scores, while the right panel displays results using unique peptides from Unipept. Correct taxonomic identifications are indicated by darker shades of blue.

**Figure 5:**
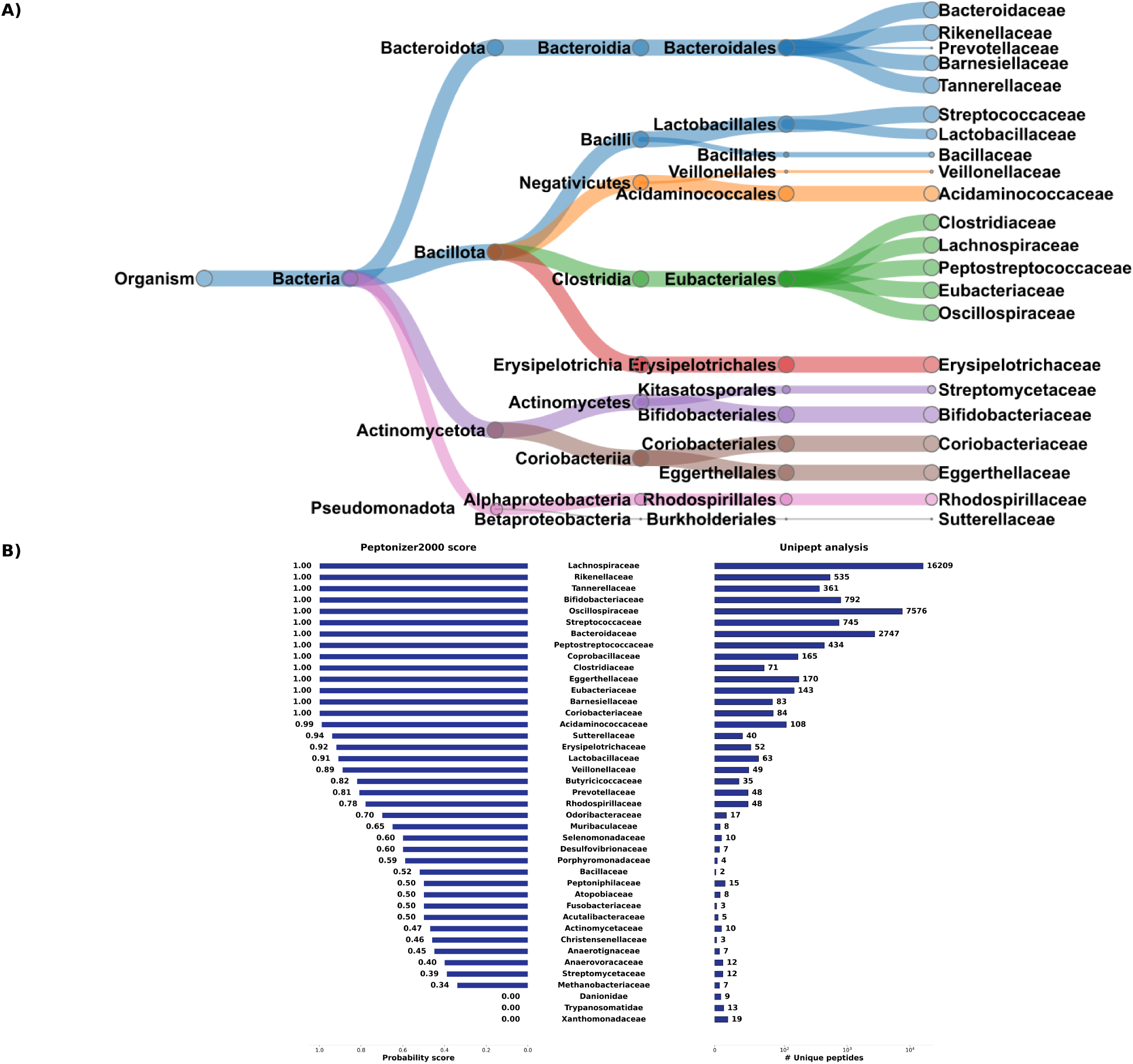
(A) Taxonomic tree view of the Peptonizer2000 results for fecal sample F07 from the CAMPI study, filtered with a probability score above 0.6. Node sizes are proportional to the probabilities attributed by the Peptonizer2000. (B) Bar plots without filtering of results showing the Peptonizer2000 analysis (left) and Unipept analysis (right).

#### Search results using untailored reference databases

To investigate the performance of the Peptonizer2000 with untailored reference databases, we searched the SIHUMIx sample S07 against two additional proteome reference databases.

First, we used the Integrated Gene Catalogue (IGC) [https://www.nature.com/articles/nbt.2942] from the CAMPI PRIDE repository. This catalog includes reference proteomes for many genera commonly found in the human gut, including those present in the SIHUMIx sample, but may not have exact species matches.

The results, presented in Figure 4, also include species-level taxonomic identifications based on unique peptides generated by Unipept. Three species present in low abundance in SIHUMIx—*Lactoplantibacillus plantarum*, *Clostridium butyricum*, and *Bifidobacterium longum*—were not detected by either Peptonizer2000 or Unipept. The use of a larger and less specific database resulted in no unique peptides being identified for these species. Consequently, the number of peptide hits was too low to be included in the Peptonizer2000 graphical model. For the five detected taxa, Peptonizer2000’s probability scores and the number of unique peptides detected allowed accurate conclusions about the sample’s composition. All five correct taxa received the highest scores with Peptonizer2000. The next highest scoring taxon, *Blautia obeum*, did not appear in the top taxa of the Unipept analysis and can thus be excluded. Relying solely on Unipept would have made it more difficult to determine which taxa were truly present.

To simulate a scenario without prior knowledge of the microbial sample, we analyzed sample S07 from CAMPI using UniRef50.^38^ With over 66 million entries as of September 2024, this large and unspecialized database significantly reduces the peptide identification rate, resulting in lower accuracy for taxonomic assignments. We also conducted an analysis using a 5% peptide FDR, in addition to the standard 1%, to see if it could improve taxonomic identification (Supplementary Figures 8–11). As anticipated, using such an extensive database impeded accurate taxonomic identification. For SIHUMIx, a maximum of four correct taxa (*Bacteroides thetaiotaomicron*, *Anaerostipes Caccae*, *Thomasclavelia Ramosa*, *Blautia producta*) appear in the top 20 results. While Peptonizer2000 showed a slight improvement with a 5% FDR, the performance of Unipept declined. This is because Peptonizer2000 considers peptide scores, allowing it to include lower-confidence peptides in the taxonomic identification while appropriately reflecting their uncertainty in the probability scores. Overall, no clear taxonomic attributions can be made using UNIREF50.

#### Analysis of a fecal sample

To evaluate the performance of the Peptonizer2000 on real microbiomes, we analyzed the fecal sample F07 from the CAMPI study. We used the IGC gut reference database. We evaluated the data at three taxonomic resolution levels: species, genus, and family. Figure 5A shows the Peptonizer2000 results at the family level with a probability score above 0.6, detecting 22 bacterial families. Overall, these results align with previous analyses of fecal samples in the CAMPI study^31^ and other human gut microbiome studies,^39,40^ demonstrating the ability of Peptonizer2000 to compute meaningful taxonomic probability scores at higher taxonomic levels. The range of the number of unique peptides per family is very large - from 1 to 16209. All families with a very large number of unique peptides are also attributed very high probability scores by the Peptonizer2000. For families with lower numbers of taxonomic attributions, the Peptonizer2000 scores can hint towards the certainty of their presence. For example, the family *Suterellaceae* has 40 unique peptides detected and a probability of presence of 0.94, and is therefore more likely to be present, which overlaps with previous analyses of this sample.^31^ *Bacillaceae* obtain a score of 0.52, even though only two unique peptides are detected. As this family of bacteria has been isolated from fecal samples before,^41^ they might truly be present in the fecal sample F07. Still, no further conclusions can be drawn since no ground truth is available. We also examined the results from Peptonizer2000 and Unipept at both genus and species levels. Although the Peptonizer2000 identified many species as present (Supplementary Figure 8), human gut microbiomes are rarely analyzed with comprehensive species-level resolution. Therefore, we focus on discussing the genus-level results shown in Figure 6. Here, the taxa assigned a high number of unique peptides by Unipept also received high Peptonizer2000 scores.

**Figure 6:**
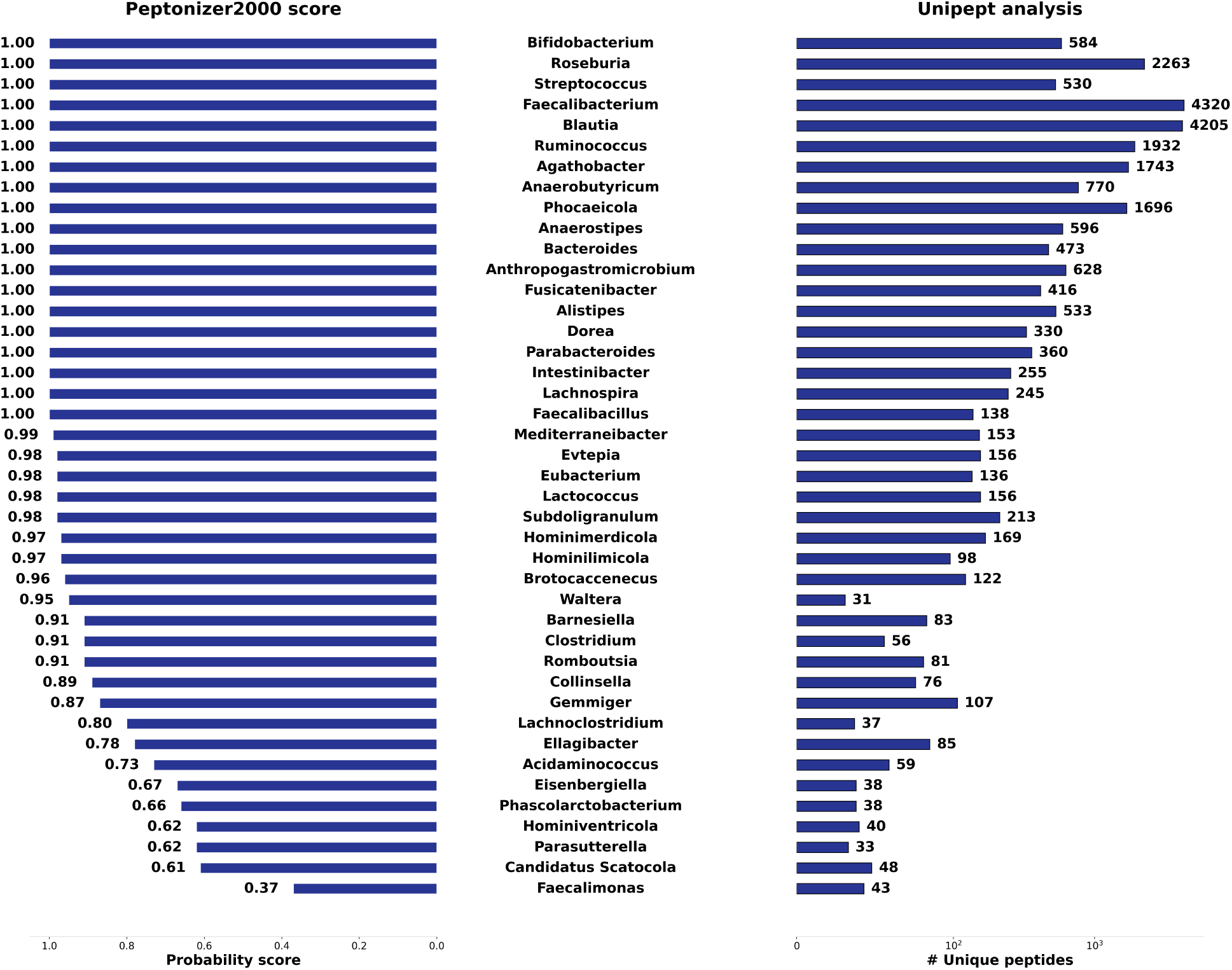
Taxonomic identification of the fecal sample F07 at the genus level. The left panel shows results using the Peptonizer2000 with corresponding probability scores, while the right panel displays results using unique peptides from Unipept.

The Unipept and Peptonizer2000 results show a high overlap. While the number of unique peptides from Unipept has a very high range ( from 33 to 4320) the Unipept scores are more evenly distributed and can ease interpretation. Notably, all genera with high scores are know to be present in the human gut. Examples include *Anaerobutyricum* (770 unique peptides, score of 1.0),^42^ *Fusicatenibacter* (416 unique peptide, score of 1),^43^ *Waltera* (33 unique peptides, score of 0.95).^44^ Again, no further conclusions can be drawn since no ground truth is available.

### Parameter choices for the graphical model

For all analyses, the Peptonizer2000 was run for the same ranges of parameters *α*, *γ*, and *β*: *α* ∈ [0.7, 0.8, 0.9, 0.99], *β* ∈ [0.2, 0.3, 0.4, 0.5, 0.6, 0.7, 0.8, 0.9], and *γ* ∈ [0.1, 0.3, 0.5]. A detailed description of these parameters is found in the Methods section. A grid search then identifies the optimal parameter set for each sample. The parameter set selected for each sample is shown in Table 1. The parameter *α* represents the probability of detecting a peptide if the parent taxon is present. Given the large number of different peptides typically detected in metaproteomics, it makes sense that *α* ≥ 0.9 was selected for all samples. If a parent taxon is present, a peptide corresponding to it will likely be observed. This parameter choice aligns with parameter ranges identified as optimal in other studies using similar graphical models. For instance, for viruses, where the probability of detecting peptides is relatively low, the optimal *α* was found to be small.^29^ For more general protein inference, *α* was identified to be much higher.^45^

**Table 1:** Parameters of the noisy OR conditional probability tables, *α*, *β*, and *γ*, selected as best fitting for each sample.

| Sample | Origin/Composition | Reference database | $\alpha$ | $\beta$ | $\gamma$ |
| --- | --- | --- | --- | --- | --- |
| s03 | CAMPI/SIHUMIx | tailored | 0.99 | 0.7 | 0.5 |
| s05 | CAMPI/SIHUMIx | tailored | 0.99 | 0.7 | 0.5 |
| s07 | CAMPI/SIHUMIx | tailored | 0.99 | 0.7 | 0.5 |
| s07 | CAMPI/SIHUMIx | IGC gut | 0.99 | 0.9 | 0.1 |
| s07 | CAMPI/SIHUMIx | UNIRef50 | 0.99 | 0.7 | 0.3 |
| s08 | CAMPI/SIHUMIx | tailored | 0.99 | 0.6 | 0.5 |
| s11 | CAMPI/SIHUMIx | tailored | 0.99 | 0.6 | 0.5 |
| s11 | CAMPI/SIHUMIx | tailored | 0.99 | 0.6 | 0.5 |
| F07 | CAMPI/fecal | IGC gut | 0.99 | 0.6 | 0.5 |
| U1 | Kleiner/28Mix | tailored | 0.99 | 0.7 | 0.5 |
| P1 | Kleiner/28Mix | tailored | 0.9 | 0.5 | 0.1 |
| C1 | Kleiner/28Mix | tailored | 0.99 | 0.7 | 0.1 |

**Table 2:** Benchmark of runtime and memory use of the Peptonizer workflow, across all parameters ranges, for samples producing graphs of different sizes.

| Sample | # nodes (N) # edges (e) | CPU time (s) min - max | Memory (GB) min - max |
| --- | --- | --- | --- |
| S03 | N = 48115, e = 197436 | 940 - 1453 | 2.5 - 2.6 |
| S05 | N = 54622, e = 239145 | 1612 - 2832 | 3.7 - 3.8 |
| S11 | N = 74434, e = 323624 | 2177 - 3426 | 4.9 - 5.1 |
| F07 | N = 30000, e = 111775 | 334 - 489 | 0.92 - 0.98 |
| U1 | N= 66289 , e = 230546 | 581 - 916 | 1.8 -1.9 |

The parameter *β* represents the probability of detecting a peptide at random but can also be interpreted as the probability for a connection between a parent taxon and a detected peptide to be erroneously present. *β* ≥ 0.5 was identified for all samples, meaning the model assumes a high probability of a wrong peptide or a wrong connection between the peptide and its parent taxon. This observation agrees with the setup of our metaproteomic identification workflow: the peptides are queried against the whole taxonomy, and in the default settings, at least 150 taxa are included in the graphical model. Protein sequence homology, especially among housekeeping proteins,^46^ but also between closely related species, contributes to this high error probability. Many species and their connections to peptides not present in the sample will, therefore, still be present in the graphical model, which is reflected in the high error probability *β*. The parameter identified for *γ* varies: *γ* = 0.5 indicates no prior knowledge of taxa presence, which is common in metaproteomics. A lower *γ* suggests that when more than 150 species are included in the graph but only 30 species are actually present in a sample, most taxa in the graphical model are more likely to be absent than present.

Reducing the number of parameter combinations can greatly reduce the computational demands of the Peptonizer2000 workflow. Based on the observed parameters for the tested samples, we recommend including the following in the grid search for future metaproteomic analyses: *α* ∈ [0.85, 0.9, 0.99], *β* ∈ [0.5, 0.6, 0.7], and *γ* ∈ [0.1, 0.3, 0.5]. These choices represent reasonable defaults and reduce the grid search space to 27 parameter combinations.

### Memory use and runtime

The Peptonizer2000 workflow consists of several steps, with the belief propagation being the most time-consuming. As of Unipept 5.0, the query time in Unipept is reduced to several minutes. The belief propagation algorithm runs until approximate convergence, and all parameter combinations can be run fully parallelized on multiple CPUs using Snakemake. If parameter sets need to be run sequentially, runtime will increase. The speed of convergence depends on the graph size, and on the distance of the initial beliefs of the graph and the final probability distribution. The worst case time complexity of the initialization of the messages for the belief propagation method is *O*(*n*^2^), where *n* is the number of nodes in the graph. We benchmarked execution times using Intel Xeon E5 - 2650 v2 CPUs for different test samples, representing different graph sizes. To take full advantage of parallelization, we recommend running the Peptonizer2000 in server environments.

## Discussion and outlook

In this study, we introduced the Peptonizer2000, a graphical model-based workflow designed to determine the taxonomic composition of microbiome samples through metaproteomic analysis. The Peptonizer2000 is implemented as a Snakemake workflow, including all analysis steps from database search to results visualization. While the current implementation includes X!Tandem^25^ as the default database search algorithm, our workflow is flexible and can integrate other database search engines as input. The primary input can also be a file with one column for detected peptides and another for attributed scores. In contrast to previous methods for metaproteomic analysis, the Peptonizer2000 incorporates peptides and their scores from database searches into the workflow, using the scores to refine taxonomic assignments. Peptide data is processed through a newly developed Unipept API endpoint, which improves the taxon inference process. Using the input, Peptonizer2000 builds a graphical model of the joint probability distribution of peptides and associated taxa. The model is subsequently calibrated, and marginal probabilities for the presence of each taxon included in the graph are computed, offering a probabilistic estimate of taxonomic composition.

We have shown that the Peptonizer2000 generates reliable probability scores for metaproteomic samples of varying complexity. In particular, we highlighted the benefit of using the probability scores together with the number of unique peptides per taxon, as identified by Unipept, to refine the determination of taxa presence or absence. This can especially help to justify the imposition of thresholds such as ’at least two unique peptides per taxon’, a heuristic very commonly used in metaproteomics. In addition, we showed that Peptonizer2000 can identify taxa, such as genera, with reasonable probability scores, even when no unique peptides are detected. This is achieved by integrating shared peptides into the graphical model, as demonstrated in a fecal microbiome sample analysis. This ability to infer the presence of taxa despite the absence of unique peptides highlights Peptonizer2000’s potential to improve the accuracy of taxonomic profiling in complex microbial communities.

The analysis of more complex lab-assembled mixtures showed that the Peptonizer2000 can compute robust probability assignments for most species present. It may not always be able to resolve closely related species within the same genus. For instance, it detected two neighboring species from the genus *Rhizobium* with high probability scores when only one was truly present. Including the number of unique peptides attributed by Unipept in the decision-making process can help resolve potential ambiguities.

As the Peptonizer2000 computes a probability distribution, imposing an exact threshold posterior probability for determining the presence or absence of a taxon is not justifiable. However, based on the previously described results, we can estimate approximate threshold probabilities. For low-complexity samples, such as SIHUMIx, the threshold should be above a score of 0.9. For medium-complexity samples, such as lab-assembled mixtures or samples analyzed with untailored but not generalist reference databases (e.g., SIHUMIx searched against IGC), the threshold should be around 0.8. For high-complexity samples or those analyzed with generalist reference databases (e.g., fecal samples searched against UniProtKB), we would not recommend any specific threshold value. The variability in sample complexity and the chosen database significantly impact the determination of appropriate threshold values, highlighting the need for careful consideration in each specific context. Alternatively, future developments in metaproteomic analysis could evolve from the currently accepted practices of setting false-discovery rates and arbitrary cut-offs towards more sensitive data analysis methods able to handle the unavoidable uncertainties of any data acquisition method and produce probabilistic error estimates.^47^

Currently, we recommend running the Peptonizer2000 in server environments due to its computational demands. In the future, several algorithmic improvements will significantly speed up the workflow.

For computing taxonomic probabilities, we currently use zero-lookahead belief propagation, which has been shown to be up to five times faster than standard belief propagation.^48^ Despite this improvement, our method remains dependent on graph size, which practically limits the number of taxa that can be included in the graphical model. To address this limitation, graph clustering approaches through community detection^49^ could be employed in the future. These approaches substantially reduce graph size and, consequently, computation time by breaking up the graph of peptides and taxa into smaller, densely connected communities. Additionally, this would enable fully parallelized computation of taxon posteriors for each community, further enhancing efficiency.

To streamline access to Peptonizer2000 for researchers, we plan to integrate it into the Unipept web application (https://unipept.ugent.be/). This integration will enhance the existing capabilities for taxonomic analysis based on unique peptides while also providing a more user-friendly web interface.

It is important to note that Peptonizer2000 relies heavily on the peptides detected through MS/MS experiments and downstream bioinformatic analyses. Ongoing research and advancements in peptide detection, such as data-independent acquisition methods for metaproteomics^19,27^ and the development of database search engines capable of effectively resolving chimeric spectra,^50^ can be expected to translate into more accurate and robust taxonomic identifications.

Looking ahead, these advancements will not only improve the speed and scalability of the Peptonizer2000 but also broaden its applicability to more complex and diverse datasets within the metaproteomics community. This progress will enable researchers to gain deeper insights from their data, ultimately advancing the understanding of microbial communities, including the identification and functional roles of their key players.

## Methods

### The Peptonizer2000 workflow

The Peptonizer2000 is a comprehensive workflow designed to analyze metaproteomics data. The workflow starts with an input file provided by the user, which contains the results of a metaproteomics sample analysis. This input file uses a generic tabular format, with one column listing peptides and a second column providing a corresponding score for each peptide. These scores represent the probability of each peptide to truly be present in the sample, as, for instance, reported by the search engine. Note that this flexible input format allows the user to apply the Peptonizer2000 workflow to search results from both DDA and DIA methodologies. Additionally, even peptides resulting from *de novo* sequencing are supported.

### (1) Unipept query for taxonomic annotation of peptides

The workflow starts by querying the Unipept database to retrieve taxonomic annotations for each peptide. All peptides provided are submitted to Unipept through a newly developed API endpoint. This API returns all taxa associated with a peptide based on its UniProt^23^ annotation without computing the LCA taxon, as was previously the case. The endpoint provides a list of all species and strains whose proteomes contain the queried peptide.

Peptides mapping to more than 10,000 proteins are excluded from the search. Peptides associated with that many proteins are likely part of so-called housekeeping proteins,^51^ which contribute minimal taxonomic information. A preliminary investigation into the LCA of the excluded proteins is shown in the supplementary materials, specifically in the section ’Taxonomic rank of peptides and proteins excluded by the Peptonizer2000 and confirms this observation.

Notably, the taxonomic range of the query can be refined by specifying a set of species, enabling users to limit the search based on prior knowledge about potentially present microorganisms in the sample. Users can restrict the taxa of interest at any taxonomic level by specifying their corresponding taxid (NCBI unique taxon identifier). For example, if the specified taxid is 2, the Unipept query will be limited to bacterial reference proteomes. In all our analyses, we did not restrict the taxonomy of the Unipept query. This approach simulates a scenario where researchers lack prior knowledge about the microbial composition of the sample.

The querying process results in a response from Unipept in JSON format, containing taxonomic information linked to the peptides, their scores, and the number of PSMs. This output is saved as an intermediate file and serves as input for the following step.

#### Selection of candidate taxa

Peptides from a metaproteomic sample can map to a wide range of taxa, influenced by both sample complexity and the taxonomic range selected for querying. An unrestricted taxonomic query often results in a graph with over 1,000 potentially present organisms, leading to highly increased computation times. The number of potentially present organisms is so high because, commonly, certain peptides from protein sequence regions that are highly conserved across species are frequently detected alongside more species-specific peptides.

To handle this complexity, we restrict the number of taxa included in the graph. By default, this number is set to 150; users can customize this number through the configuration file. Every taxon with at least one unique peptide will be included in the graph, even if this results in exceeding the default limit of 150 taxa.

The number of peptides detected for a present taxon exhibits a large dynamic range. It depends both on the organism’s size and the relative abundance of the organism within the sample. For instance, in a MS run involving the commonly used, lab-assembled SIHUMIx mixture,^52^ 5 PSMs were detected for *Lactobacillus plantarum*, while over 1,000 PSMs mapped to the more abundant *Bacteroides thetaiotaomicron*. Both taxa are present in the SIHUMIx mixture. We aim to address this issue by assigning a high weight to unique PSMs. This approach makes sure that both the number of spectra matching to a peptide and the peptide’s taxonomic information, including its sequence degeneracy, are considered.

The formula to compute the weight *W_T_* of a taxon is the following:

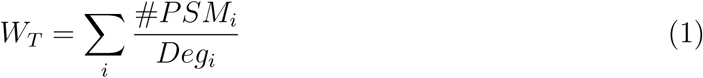

Where *i* stands for all peptides mapped to taxon T, #*PSM_i_* denotes the number of PSMs for peptide *i*, and *Deg_i_* is the degeneracy of peptide *i*—specifically, the number of taxa to which the peptide has been mapped.

All candidate taxa and their corresponding scored peptides are exported as a CSV file, which serves as the basis for the graphical model.

### (2) Assembly of the graphical model, belief propagation and grid search

#### Graphical model assembly

The construction of the graphical model, which is a factor graph, uses the same algorithm as PepGM. All peptides and taxa from step (2) are included in the graph. Two categories of nodes represent peptides and taxa, respectively. An edge is drawn between a peptide node and a taxon node if a peptide is part of a taxon’s proteome. As described in Holstein et al.,^29^ factor nodes based on the noisy-OR model and convolution tree nodes are added to the graph. Together, the graphical model represents the probability distribution of all peptides and taxa. Note that this probability distribution is an approximation, as not all taxa are included in the graph, and it relies on the same assumptions regarding peptide emission and detection probabilities as in PepGM. The noisy-OR model introduces three parameters:

- *α*: The probability of observing a peptide given the presence of its parent taxon.
- *β*: The probability of a peptide being randomly or falsely observed or incorrectly linked to a taxon that is not actually present in the sample.
- *γ*: The prior probability of a taxon being present.

We have discussed the choice and use of these parameters more extensively for PepGM.^29^ In the Peptonizer2000, we introduce an additional option to regularize the probabilities for a peptide having *n* parent taxa present in the noisy-OR model.This regularization means that the probabilities for a peptide to have *N* parent taxa present is assumed to decrease inversely proportional to *N*. This assumption has been previously applied for protein inference^45^ models probability distribution for samples where a very broad taxonomic range was queried compared to the expected number of present taxa. The user can choose to turn this regularization on or off. In metaproteomics, reference databases typically contain far more taxa than are actually present or detected in the sample. For this reason, we generally recommend keeping the regularization option activated for smaller lab-assembled mixtures. For complex biological samples, we recommend to keep the regularization option off.

#### Inference algorithm and grid search

To compute the marginal probabilities of taxa, we use the belief propagation algorithm.^53^ To efficiently handle the large size of the graph, we implemented a version of belief propagation known as ‘zero-lookahead’.^48^ This version of belief propagation was shown to scale nearly linearly with graph size while introducing only a slight approximation of the global probability distribution.

Additionally, we perform a grid search across the three model parameters introduced by the noisy-OR model. To reduce the search space, the prior probability is set to the following values: *γ* ∈ [0.1, 0.3, 0.5]. The peptide emission probability is set to *α* ∈ [0.85, 0.9, 0.99]. The error probability is set to *β* ∈ [0.5, 0.6, 0.7] by default. However, these values can be modified by the user. The rationale behind the selection of these parameters, as well as their interpretation, is discussed in the results section. The optimal parameters for a specific sample are determined in the following step.

### (3) Selection of most appropriate parameter range

For each combination of model parameters *α*, *β*, and *γ*, marginal probabilities are computed for all taxa in the graphical model. To identify the optimal set of parameters for the given sample, a thorough evaluation of these parameters is conducted.

The evaluation of the best fitting parameters is based on a comparison between a pre-defined list of weighted taxa used as input for the graphical model and the results of the belief propagation from Peptonizer2000. In the following, we refer to the weighted list as *L_W_* and the scored list as *L_S_*. Both lists of taxa are ranked according to their weights and scores, respectively.

Since the weights assigned to each taxon rely on the number and uniqueness of PSMs per taxon, the weighted taxon list *L_W_* gives an initial indication of the taxa likely to be present. However, it has one major limitation: closely related taxa, such as multiple highly similar bacterial species, may all appear near the top of the list in *L_W_*. To resolve this, we aim to select a parameter range that can distinguish between these taxa.

The solution we propose is to cluster *L_W_* based on taxon similarity. The clustering procedure is analogous to the taxonomic clustering method previously described in Alves and Yu,^24^ with one key modification: instead of relying on the full theoretical peptidome, the similarity between two taxa is based solely on their detected peptides. This adjustment ensures that the clustering is closer to the currently analyzed sample, where taxon abundances and the number of detected peptides may vary greatly.

The taxon with the highest number of detected peptides within a group of similar taxa is designated as the cluster head. A threshold similarity of 0.9 is used to group taxa into clusters—this threshold is arbitrary but was chosen to ensure that closely related taxa are clustered together. While the threshold should be well above 0.5 for meaningful clustering, a perfect similarity score of 1 is not required. In fact, slight variations in the threshold of up to 0.1 result in changes of only one or two taxa in the clustered taxon list. After taxon clustering, we obtain a simplified version of the weighted taxon list *L_W_*, which contains only the weight-sorted cluster heads.

Both lists are then ready to be compared. For the comparison, we need to take into account the following: taxa listed in the results of Peptonizer2000 *L_S_* may be absent from the clustered list *L_w_*. Thus, we need a comparison method that allows for non-conjointness. Additionally, higher-ranked taxa in both lists are more important than lower-ranked ones, meaning that taxa with higher weights and scores should have a greater influence on the comparison. To meet these criteria, we use rank-biased overlap^54^ (denoted *RBO*) as the similarity measure for list comparison. It handles non-conjointness and gives more weight to the top-ranked taxa in both lists.

The optimal parameter range for a given sample is selected by maximizing the rank-biased overlap between the clustered *L_W_* and the scored results *L_S_*while simultaneously minimizing the entropy of the marginal probability distribution. Entropy, denoted as *S*, has been previously used as a criterion in PepGM,^29^ and its inclusion here aims to maximize the information content of the resulting distribution.

Thus, the final metric that the chosen parameter range for each sample should maximize is given by:

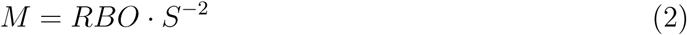

By comparing *M* across different parameter ranges, the one with the highest value can be selected, ensuring both strong alignment between *L_W_* and *L_S_* and a low-entropy, high-information distribution.

### Results output and visualization

Peptonizer2000 outputs results in various formats. The most basic output is a CSV file containing a list of all identified taxa with their corresponding score, which is the most basic output level. A bar chart summarizes the results as a .png, alongside a second bar chart that shows the performance of each parameter set based on the rank-biased-overlap and entropy metric.

A visually appealing tree view of the results can also be generated in both HTML and SVG formats, offering an interactive and high-quality graphical representation of the data.

## Materials

We evaluated Peptonizer2000 using several publicly available datasets from metaproteomic samples, which can be accessed through the PRIDE database.^55^ The first set of samples was obtained from the CAMPI study,^31^ all available under the PRIDE identifier PXD023217.

The first sample type we analyze is the SIHUMIx sample.^56^ It was developed as a model community for the human gut microbiota and covers the three dominant genera found in human feces: Firmicutes, Proteobacteria, and Bacteroidetes. It contains eight microbes: *Bacteroides thetaiotaomicron*, *Anaerostipes caccae*, *Escherischia coli*, *Lactoplantibacillus plantarum*, *Clostridium butyricum*, *Thomasclavelia ramosa*, *Blautia producta*, and *Bifidobacterium longum*. SIHUMIx is widely used for benchmarking in metaproteomics. Analogous to the original CAMPI publication, we use samples S03, S05, S07, S08, and S11 for our investigation, referring to them collectively as S03-S11. Each sample was acquired using different laboratory workflows, resulting in slightly varying coverage of the proteomes contained in the samples.

The second sample type taken from the CAMPI study is a fecal sample, for which the taxonomic ’ground truth’ is unknown. For our analysis, we chose the fecal sample denoted as F07.

Another set of samples we analyze consists of a more complex, lab-assembled mixture of up to 32 different microorganisms.^33^ These samples are available on PRIDE under the identifier PXD006118. For our analysis, we selected samples from three types of communities: those assembled with equal cell amounts per taxon (denoted ’C1’), equal protein amounts per taxon (denoted ’P1’), and uneven composition (denoted ’U1’).

For our evaluation, we used four different reference databases. The first is the SIHUMIx database, specifically tailored to the SIHUMIx sample, which is accessible through the corresponding PRIDE repository associated with the CAMPI study. The second database is the integrated gene catalog (IGC),^57^ a gut microbiota reference database that includes representative sequences aiming to cover the majority of gut microbes. The third database is tailored to the 32 potentially present microorganisms in the complex mock community and was provided alongside the experimental data of the Kleiner *et al.* study.^33^ These three databases are concatenated with a contaminants database^58^ to ensure that peptide hits matching the contaminant sequences will be filtered out before taxonomic analysis.

Finally, as a fourth reference, we use the UniREF50 database,^38^ a clustered set of sequences derived from UniProt, aiming to maintain comprehensive coverage of the sequence space while reducing redundancy.

We analyzed all samples beginning with a standard database search followed by rescoring. The MS/MS spectra are searched against the provided reference database using a combination of X!Tandem^25^ and MS2Rescore.^26^ This process generates a list of identified peptide-spectrum matches along with their corresponding scores. MS2Rescore is employed to enhance the reliability of these identifications. It rescales the initial scores based on statistical models and provides e-values that represent the probability of false detection for each peptide.

For the proteomic database search using X!Tandem, we fixed the search parameters in accordance with the original sample publications. The parameters were the same for all samples analyzed and included specific trypsin digestion with a maximum of two missed cleavages, mass tolerances of 10.0 ppm for MS1 and 0.02 Da for MS2, and fixed modifications of carbamidomethylation of cysteine (+57.021464 Da). Variable modifications included oxidation of methionine (+15.994915 Da). Additionally, we set the peptide length to range between 6 and 50 amino acids and included charge states of +2, +3, or +4. A false discovery rate (FDR) threshold of 1% was also applied to ensure the accuracy of the identified peptides. To perform the Unipept analysis, we used the Unipept command line tool with the option ’–pept2lca’, taking a .txt file containing all peptides that were provided as input for Peptonizer2000. Since Unipept outputs the LCA for the queried peptides, we filtered the Unipept response to retain only those peptides that are unique to taxa at the species level. We then aggregated the number of unique peptides per species to obtain taxonomic identifications at the species level based on unique PSMs. For analyses at higher taxonomic levels, we used the same approach, mapping unique peptides at lower taxonomic levels back to their corresponding ancestors as needed.

All code used for downstream analysis is also publicly available at https://github.com/BAMeScience/Peptonizer2000. The code used to generate the tree view graphics of the results can be found at https://github.com/pverscha/peptonizer-visualizations.

## Acknowledgements

TH was supported by the German Research Foundation (DFG) (MU 4430/2-1 to T.M.) and acknowledges additional support by the Joachim-Herz-Foundation. L.M. acknowledges funding from the Research Foundation Flanders (FWO) [G028821N] and [G010023N]. PV acknowledges funding by Ghent University [BOF/01P10623]. T.V.D.B. acknowledges funding from the Research Foundation Flanders (FWO) [1286824N].

## Notes

https://github.com/compomics/Peptonizer2000

### Competing Interest Statement

The authors have declared no competing interest.

### Summary of Updates

Updated figures, algorithm description and aper structure

## References

(1) Sunagawa, S. et al. Structure and function of the global ocean microbiome. Science 2015, 348, 1261359, Publisher: American Association for the Advancement of Science.

(2) Heyer, R.; Kohrs, F.; Reichl, U.; Benndorf, D. Metaproteomics of complex microbial communities in biogas plants. Microbial Biotechnology 2015, 8, 749–763, eprint: https://onlinelibrary.wiley.com/doi/pdf/10.1111/1751-7915.12276.

(3) Cani, P. D. Human gut microbiome: hopes, threats and promises. Gut 2018, 67, 1716– 1725, Publisher: BMJ Publishing Group Section: Recent advances in basic science.

(4) Kleiner, M. et al. Metaproteomics of a gutless marine worm and its symbiotic microbial community reveal unusual pathways for carbon and energy use. Proceedings of the National Academy of Sciences 2012, 109, E1173–E1182, Publisher: National Academy of Sciences Section: PNAS Plus.

(5) Fierer, N. Embracing the unknown: disentangling the complexities of the soil microbiome. Nature Reviews Microbiology 2017, 15, 579–590, Number: 10 Publisher: Nature Publishing Group.

(6) Pan, S.; Hullar, M. A. J.; Lai, L. A.; Peng, H.; May, D. H.; Noble, W. S.; Raftery, D.; Navarro, S. L.; Neuhouser, M. L.; Lampe, P. D.; Lampe, J. W.; Chen, R. Gut Microbial Protein Expression in Response to Dietary Patterns in a Controlled Feeding Study: A Metaproteomic Approach. Microorganisms 2020, 8, 379, Number: 3 Publisher: Multi-disciplinary Digital Publishing Institute.

(7) Shreiner, A. B.; Kao, J. Y.; Young, V. B. The gut microbiome in health and in disease. Current opinion in gastroenterology 2015, 31, 69–75.

(8) Ma, C.; Sun, Z.; Zeng, B.; Huang, S.; Zhao, J.; Zhang, Y.; Su, X.; Xu, J.; Wei, H.; Zhang, H. Cow-to-mouse fecal transplantations suggest intestinal microbiome as one cause of mastitis. Microbiome 2018, 6, 200.

(9) Wolf, M.; Schallert, K.; Knipper, L.; Sickmann, A.; Sczyrba, A.; Benndorf, D.; Heyer, R. Advances in the clinical use of metaproteomics. Expert Review of Proteomics 2023, 20, 71–86, Publisher: Taylor & Francis eprint: 10.1080/14789450.2023.2215440.

(10) Armengaud, J. Metaproteomics to understand how microbiota function: The crystal ball predicts a promising future. Environmental Microbiology 2023, 25, 115–125, eprint: https://onlinelibrary.wiley.com/doi/pdf/10.1111/1462-2920.16238.

(11) Wang, Y.; Zhou, Y.; Xiao, X.; Zheng, J.; Zhou, H. Metaproteomics: A strategy to study the taxonomy and functionality of the gut microbiota. Journal of Proteomics 2020, 219, 103737.

(12) Muth, T.; Benndorf, D.; Reichl, U.; Rapp, E.; Martens, L. Searching for a needle in a stack of needles: challenges in metaproteomics data analysis. Molecular BioSystems 2013, 9, 578–585, Publisher: The Royal Society of Chemistry.

(13) Saito, M. A. et al. Progress and Challenges in Ocean Metaproteomics and Proposed Best Practices for Data Sharing. Journal of Proteome Research 2019, 18, 1461–1476, Publisher: American Chemical Society.

(14) Schiebenhoefer, H.; Van Den Bossche, T.; Fuchs, S.; Renard, B. Y.; Muth, T.; Martens, L. Challenges and promise at the interface of metaproteomics and genomics: an overview of recent progress in metaproteogenomic data analysis. Expert Review of Proteomics 2019, 16, 375–390, Publisher: Taylor & Francis eprint: 10.1080/14789450.2019.1609944.

(15) Muth, T.; Behne, A.; Heyer, R.; Kohrs, F.; Benndorf, D.; Hoffmann, M.; Lehtevä, M.; Reichl, U.; Martens, L.; Rapp, E. The MetaProteomeAnalyzer: A Powerful Open-Source Software Suite for Metaproteomics Data Analysis and Interpretation. Journal of Proteome Research 2015, 14, 1557–1565, Publisher: American Chemical Society.

(16) Li, L.; Ning, Z.; Cheng, K.; Zhang, X.; Simopoulos, C. M. A.; Figeys, D. iMetaLab Suite: A one-stop toolset for metaproteomics. iMeta 2022, 1, e25, eprint: https://onlinelibrary.wiley.com/doi/pdf/10.1002/imt2.25.

(17) Jagtap, P. D.; Blakely, A.; Murray, K.; Stewart, S.; Kooren, J.; Johnson, J. E.; Rhodus, N. L.; Rudney, J.; Griffin, T. J. Metaproteomic analysis using the Galaxy framework. Proteomics 2015, 15, 3553–3565.

(18) Verschaffelt, P.; Van Den Bossche, T.; Martens, L.; Dawyndt, P.; Mesuere, B. Unipept Desktop: A Faster, More Powerful Metaproteomics Results Analysis Tool. Journal of Proteome Research 2021, 20, 2005–2009, Publisher: American Chemical Society.

(19) Gómez-Varela, D.; Xian, F.; Grundtner, S.; Sondermann, J. R.; Carta, G.; Schmidt, M. Increasing taxonomic and functional characterization of host-microbiome interactions by DIA-PASEF metaproteomics. Frontiers in Microbiology 2023, 14, Publisher: Frontiers.

(20) Gouveia, D.; Pible, O.; Culotta, K.; Jouffret, V.; Geffard, O.; Chaumot, A.; Degli-Esposti, D.; Armengaud, J. Combining proteogenomics and metaproteomics for deep taxonomic and functional characterization of microbiomes from a non-sequenced host. npj Biofilms and Microbiomes 2020, 6, 1–6, Publisher: Nature Publishing Group.

(21) Bihani, S.; Gupta, A.; Mehta, S.; Rajczewski, A. T.; Johnson, J.; Borishetty, D.; Griffin, T. J.; Srivastava, S.; Jagtap, P. D. Metaproteomic Analysis of Nasopharyngeal Swab Samples to Identify Microbial Peptides in COVID-19 Patients. Journal of Proteome Research 2023, 22, 2608–2619, Publisher: American Chemical Society.

(22) Bender, M. A.; Farach-Colton, M.; Pemmasani, G.; Skiena, S.; Sumazin, P. Lowest common ancestors in trees and directed acyclic graphs. Journal of Algorithms 2005, 57, 75–94.

(23) UniProt: a hub for protein information. Nucleic Acids Research 2015, 43, D204–D212.

(24) Alves, G.; Yu, Y.-K. Robust Accurate Identification and Biomass Estimates of Microorganisms via Tandem Mass Spectrometry. Journal of the American Society for Mass Spectrometry 2020, 31, 85–102.

(25) Craig, R.; Beavis, R. C. TANDEM: matching proteins with tandem mass spectra. Bioinformatics 2004, 20, 1466–1467.

(26) Declercq, A.; Bouwmeester, R.; Hirschler, A.; Carapito, C.; Degroeve, S.; Martens, L.; Gabriels, R. MS2Rescore: Data-Driven Rescoring Dramatically Boosts Immunopeptide Identification Rates. Molecular & Cellular Proteomics 2022, 21, 100266.

(27) Pietilä, S.; Suomi, T.; Elo, L. L. Introducing untargeted data-independent acquisition for metaproteomics of complex microbial samples. ISME Communications 2022, 2, 51.

(28) Rajczewski, A. T.; Blakeley-Ruiz, J. A.; Meyer, A.; Vintila, S.; McIlvin, M. R.; Bossche, T. V. D.; Searle, B. C.; Griffin, T. J.; Saito, M. A.; Kleiner, M.; Jagtap, P. D. Data-Independent Acquisition Mass Spectrometry as a Tool for Metaproteomics: Inter- laboratory Comparison Using a Model Microbiome. 2024; https://www.biorxiv.org/content/10.1101/2024.09.18.613707v1, Pages: 2024.09.18.613707 Section: New Results.

(29) Holstein, T.; Kistner, F.; Martens, L.; Muth, T. PepGM: a probabilistic graphical model for taxonomic inference of viral proteome samples with associated confidence scores. Bioinformatics 2023, 39, btad289.

(30) Mölder, F.; Jablonski, K. P.; Letcher, B.; Hall, M. B.; Tomkins-Tinch, C. H.; Sochat, V.; Forster, J.; Lee, S.; Twardziok, S. O.; Kanitz, A.; Wilm, A.; Holtgrewe, M.; Rahmann, S.; Nahnsen, S.; Köster, J. Sustainable data analysis with Snakemake. F1000Research 2021, 10, 33.

(31) Van Den Bossche, T., et al. Critical Assessment of MetaProteome Investigation (CAMPI): a multi-laboratory comparison of established workflows. Nature Communications 2021, 12, 7305, Number: 1 Publisher: Nature Publishing Group.

(32) Van Den Bossche, T., et al. The Metaproteomics Initiative: a coordinated approach for propelling the functional characterization of microbiomes. Microbiome 2021, 9, 243.

(33) Kleiner, M.; Thorson, E.; Sharp, C. E.; Dong, X.; Liu, D.; Li, C.; Strous, M. Assessing species biomass contributions in microbial communities via metaproteomics. Nature Communications 2017, 8, 1558, Number: 1 Publisher: Nature Publishing Group.

(34) Yan, P.; Sun, Y.; Luo, J.; Liu, X.; Wu, J.; Miao, Y. Integrating the serum proteomic and fecal metaproteomic to analyze the impacts of overweight/obesity on IBD: a pilot investigation. Clinical Proteomics 2023, 20, 6.

(35) Pettersen, V. K.; Dufour, A.; Arrieta, M.-C. Metaproteomic profiling of fungal gut colonization in gnotobiotic mice. Animal Microbiome 2022, 4, 14.

(36) Oren, A. Nomenclature of prokaryotic ‘Candidatus’ taxa: establishing order in the current chaos. New Microbes and New Infections 2021, 44, 100932.

(37) Armengaud, J.; Trapp, J.; Pible, O.; Geffard, O.; Chaumot, A.; Hartmann, E. M. Non-model organisms, a species endangered by proteogenomics. Journal of Proteomics 2014, 105, 5–18.

(38) Suzek, B. E.; Wang, Y.; Huang, H.; McGarvey, P. B.; Wu, C. H. UniRef clusters: a comprehensive and scalable alternative for improving sequence similarity searches. Bioinformatics 2015, 31, 926–932.

(39) Brown, M. D.; Shinn, L. M.; Reeser, G.; Browning, M.; Schwingel, A.; Khan, N. A.; Holscher, H. D. Fecal and soil microbiota composition of gardening and non-gardening families. Scientific Reports 2022, 12, 1595, Publisher: Nature Publishing Group.

(40) Almeida, A.; Nayfach, S.; Boland, M.; Strozzi, F.; Beracochea, M.; Shi, Z. J.; Pollard, K. S.; Sakharova, E.; Parks, D. H.; Hugenholtz, P.; Segata, N.; Kyrpides, N. C.; Finn, R. D. A unified catalog of 204,938 reference genomes from the human gut microbiome. Nature Biotechnology 2021, 39, 105–114, Publisher: Nature Publishing Group.

(41) Hoyles, L.; Honda, H.; Logan, N. A.; Halket, G.; La Ragione, R. M.; McCartney, A. L. Recognition of greater diversity of *Bacillus* species and related bacteria in human faeces. Research in Microbiology 2012, 163, 3–13.

(42) Ramirez Garcia, A.; Greppi, A.; Constancias, F.; Ruscheweyh, H.-J.; Gasser, J.; Hurley, K.; Sturla, S. J.; Schwab, C.; Lacroix, C. Anaerobutyricum hallii promotes the functional depletion of a food carcinogen in diverse healthy fecal microbiota. Frontiers in Microbiomes 2023, 2, Publisher: Frontiers.

(43) Zhou, J.; Wu, X.; Li, Z.; Zou, Z.; Dou, S.; Li, G.; Yan, F.; Chen, B.; Li, Y. Alterations in Gut Microbiota Are Correlated With Serum Metabolites in Patients With Insomnia Disorder. Frontiers in Cellular and Infection Microbiology 2022, 12, Publisher: Frontiers.

(44) Liu, C. et al. Enlightening the taxonomy darkness of human gut microbiomes with a cultured biobank. Microbiome 2021, 9, 119.

(45) Pfeuffer, J.; Sachsenberg, T.; Dijkstra, T. M. H.; Serang, O.; Reinert, K.; Kohlbacher, O. EPIFANY: A Method for Efficient High-Confidence Protein Inference. Journal of Proteome Research 2020, 19, 1060–1072, Publisher: American Chemical Society.

(46) https://orcid.org/0000 0002-9339-2511, C. J. J.; https://orcid.org/0000 0002-7047-5445, W. K.; Drangowska-Way, A.; https://orcid.org/0000 0003-0503-4181, E. J. O.; https://orcid.org/0000 0001-7700-3654, N. E. L. What are housekeeping genes? PLoS Computational Biology 2022, 18, Place: San Francisco, United States Publisher: Public Library of Science Section: Research Article.

(47) The, M.; Käll, L. Integrated Identification and Quantification Error Probabilities for Shotgun Proteomics * [S]. Molecular & Cellular Proteomics 2019, 18, 561–570.

(48) Sutton, C.; McCallum, A. Improved dynamic schedules for belief propagation. Proceedings of the Twenty-Third Conference on Uncertainty in Artificial Intelligence. Arlington, Virginia, USA, 2007; pp 376–383.

(49) Lancichinetti, A.; Fortunato, S. Community detection algorithms: A comparative analysis. Physical Review E 2009, 80, 056117, Publisher: American Physical Society.

(50) Dorfer, V.; Maltsev, S.; Winkler, S.; Mechtler, K. CharmeRT: Boosting Peptide Identifications by Chimeric Spectra Identification and Retention Time Prediction. Journal of Proteome Research 2018, 17, 2581–2589.

(51) Joshi, C. J.; Ke, W.; Drangowska-Way, A.; O’Rourke, E. J.; Lewis, N. E. What are housekeeping genes? PLOS Computational Biology 2022, 18, e1010295, Publisher: Public Library of Science.

(52) Krause, J. L.; Schaepe, S. S.; Fritz-Wallace, K.; Engelmann, B.; Rolle-Kampczyk, U.; Kleinsteuber, S.; Schattenberg, F.; Liu, Z.; Mueller, S.; Jehmlich, N.; Von Bergen, M.; Herberth, G. Following the community development of SIHUMIx – a new intestinal in vitro model for bioreactor use. Gut Microbes 2020, 11, 1116–1129, Publisher: Taylor & Francis eprint: 10.1080/19490976.2019.1702431.

(53) Knoll, C.; Rath, M.; Tschiatschek, S.; Pernkopf, F. Message Scheduling Methods for Belief Propagation. Machine Learning and Knowledge Discovery in Databases. Cham, 2015; pp 295–310.

(54) Webber, W.; Moffat, A.; Zobel, J. A similarity measure for indefinite rankings. ACM Transactions on Information Systems 2010, 28, 20:1–20:38.

(55) PRIDE: The proteomics identifications database – Martens - 2005 - PROTEOMICS - Wiley Online Library. https://analyticalsciencejournals.onlinelibrary.wiley.com/doi/abs/10.1002/pmic.200401303?casa_token=HwpGOoawKj4AAAAA%3AMjh_J-lycbwXZNY-R3ESEcl4-UTWNFNrkLMKt6ht5toZwcGt9ht9lzWC2-rBvgzlySrtiNe_tcKWFg.

(56) Schäpe, S. S.; Krause, J. L.; Engelmann, B.; Fritz-Wallace, K.; Schattenberg, F.; Liu, Z.; Müller, S.; Jehmlich, N.; Rolle-Kampczyk, U.; Herberth, G.; von Bergen, M. The Simplified Human Intestinal Microbiota (SIHUMIx) Shows High Structural and Functional Resistance against Changing Transit Times in In Vitro Bioreactors. Microorganisms 2019, 7, 641.

(57) Li, J. et al. An integrated catalog of reference genes in the human gut microbiome. Nature Biotechnology 2014, 32, 834–841, Publisher: Nature Publishing Group.

(58) Frankenfield, A. M.; Ni, J.; Ahmed, M.; Hao, L. Protein Contaminants Matter: Building Universal Protein Contaminant Libraries for DDA and DIA Proteomics. Journal of Proteome Research 2022, 21, 2104–2113, Publisher: American Chemical Society.

