## Supplementary_materials for "The Peptonizer2000: bringing confidence to metaproteomics"

### Additional information regarding the Peptonizer2000 workflow

#### List of packages used

Table 1: Packages used in the Peptonizer2000

| Package name | reference |
|---|---|
| Ete3 | [1] |
| pandas | [2] |
| numpy | [?] |
| networkx | [3] |
| scipy | [4] |
| Biopython | [5] |
| matplotlib | [6] |
| seaborn | [7] |

### Taxonomic rank of peptides and proteins excluded by the Peptonizer

The Peptonizer2000 excludes peptides from its Unipept query that map to more than 10000 proteins. In the following, we describe which peptide taxon mappings they exclude and why this leads to no, or only a negligible reduction in taxonomic resolution.

Every peptide sequence in the Unipept database is associated with one or more proteins in which the sequence occurs. The amount of proteins that a peptide is associated with can differ drastically. The following table presents an overview of how many peptides are associated with $n$ or more proteins:

Table 2 shows that the amount of peptides associated with a large number of proteins decreases rapidly. There are only a little over 13000 peptides that occur in 10000 or more proteins. For these peptides, we expect the lowest common ancestor to be very generic, as they might, e.g., originate from housekeeping proteins. In the following, we analyse how far

Table 2: number of peptides associated to $n$ or more Proteins in the Unipept database

| # associated proteins $n$ | # peptides |
|---|---|
| $\geq 1$ | 1 342 470 764 |
| $\geq 2$ | 355 979 324 |
| $\geq 10$ | 38 697 210 |
| $\geq 10^2$ | 2 921 879 |
| $\geq 10^3$ | 217 922 |
| $\geq 10^4$ | 13 008 |
| $\geq 10^5$ | 118 |
| $\geq 10^6$ | 0 |

this expectation holds.

The 13000 peptides mapping to 10000 or more proteins are extracted and queried for their LCA. The results are shown in table 4.

Indeed, about 94% of the excluded peptides have 'root' as their LCA and therefore carry no taxonomic information. It is therefore sound to exclude these peptides from the Peptonizer analysis.

Interestingly, there are the 200 peptide sequences with an LCA at the species rank. Table 4 shows which species the peptide with more than 10000 proteins associated, but with high taxon specificity, map to.

The majority of these species are viral. For most viruses shown, such as HIV or influenza, research is intensive and enormous amounts of different strains exist. Many referneces are uploaded to the Uniprot database by scientist, explaining why there are 200 peptides that occur in 10000 or more proteins and that still have an LCA annotated at the species rank. On a side note, this showcases he bias towards model organisms and certain species or strains in public proteome databases.

Table 3: LCA of the 13000 peptides mapping to more than 10000 proteins.

| NCBI taxonomy rank | Peptides |
|---|---|
| root | 12 369 |
| superkingdom | 43 |
| kingdom | 16 |
| subkingdom | 0 |
| superphylum | 0 |
| phylum | 8 |
| subphylum | 7 |
| superclass | 1 |
| class | 18 |
| subclass | 1 |
| superorder | 0 |
| order | 0 |
| infraorder | 1 |
| superfamily | 0 |
| family | 2 |
| subfamily | 0 |
| tribe | 1 |
| subtribe | 0 |
| genus | 55 |
| subgenus | 0 |
| species group | 0 |
| species subgroup | 0 |
| species | 200 |
| subspecies | 0 |
| strain | 1 |
| varietas | 0 |
| forma | 0 |

Table 4: Amount of species-specific peptides per species that map to more than 10000 proteins.

| # Peptides | LCA |
|---|---|
| 119 | Alphainfluenzavirus influenzae |
| 32 | Human immunodeficiency virus |
| 14 | Hepatitis B virus |
| 9 | Betainfluenzavirus influenzae |
| 4 | Orthoflavivirus denguei |
| 3 | Simian immunodeficiency virus |
| 1 | Alcidodes juglans |
| 1 | Bacillus subtilis |
| 1 | Bacteroides thetaiotaomicron |
| 1 | Cannabis sativa |
| 1 | Capsicum baccatum |
| 1 | Echinocucumis hispida |
| 1 | Geissoloma marginatum |
| 1 | Homo sapiens |
| 1 | Human immunodeficiency virus |
| 1 | Kalanchoe fedtschenkoi |
| 1 | Leucosceptrum canum |
| 1 | Loxia curvirostra |
| 1 | Marinilactibacillus piezotolerans |
| 1 | Melanocenchris jacquemontii |
| 1 | Merops nubicus |
| 1 | Morbillivirus hominis |
| 1 | Phalaenopsis pulcherrima |
| 1 | Phormidesmis priestleyi |
| 1 | Rhodobacter maris |

### Composition of the samples C1, P1, U1

The composition of the lab assembles samples C1,P1 und U1 was described in.[8] For convenience, we resume the composition of the three samples in the table below.

Table 5: Taxonomic composition of the three lab assembled communities U, C and P. Units are in $\mu g$

| **Species** | U | C | P |
|---|---|---|---|
| Agrobacterium tumefaciens | 4186.28 | 361.51 | 642.7 |
| Alteromonas macleodii | 707.47 | 214.38 | 642.7 |
| Bacillus subtilis | 583.83 | 717.06 | 642.7 |
| Burkholderia xenovorans | 321.37 | 69.86 | 642.7 |
| Chlamydomonas reinhardtii | 2962.73 | 6583.85 | 642.7 |
| Chromobacterium violaceum | 933.08 | 741.72 | 642.7 |
| Cupriavidus metallidurans | 11504.98 | 216.26 | 642.7 |
| Desulfovibrio vulgaris | 701.58 | 0.00 | 0 |
| Escherichia coli | 4290.54 | 319.24 | 642.7 |
| Nitrosomonas europaeae | 60.58 | 0.00 | 0 |
| Nitrosomonas ureae | 402.68 | 0.00 | 0 |
| Nitrososphaera viennensis | 607.41 | 27.61 | 642.7 |
| Nitrosospira multiformis | 155.16 | 0.00 | 0 |
| Paracoccus denitrificans | 683.66 | 198.74 | 642.7 |
| Phage ES18 | 65.55 | 45.20 | 62 |
| Phage F0 | 65.28 | 5.58 | 62 |
| Phage F2 | 62.04 | 62.04 | 62 |
| Phage M13 | 109.02 | 109.02 | 62 |
| Phage P22 | 78.77 | 0.93 | 62 |
| Pseudomonas denitrificans | 2128.51 | 551.43 | 642.7 |
| Pseudomonas fluorescens | 4964.22 | 738.72 | 642.7 |
| Pseudomonas pseudoalcaligenes | 863.80 | 716.25 | 642.7 |
| Rhizobium leguminosarum bv. viciae 3841 | 680.47 | 348.96 | 642.7 |
| Rhizobium leguminosarum bv. viciae VF39 | 1671.03 | 185.67 | 642.7 |
| Roseobacter sp. AK199 | 1183.43 | 408.08 | 642.7 |
| Salmonella enterica typhimurium (3 strains combined) | 25037.78 | 1123.11 | 1928.1 |
| Staphylococcus aureus ATCC 13709 | 715.15 | 30.05 | 642.7 |
| Staphylococcus aureus ATCC 25923 | 1216.31 | 24.04 | 642.7 |
| Stenotrophomonas maltophilia | 5946.27 | 448.44 | 642.7 |
| Thermus Thermophilus | 1245.63 | 386.84 | 642.7 |

### Additional Results plots

#### SIHUMIx samples

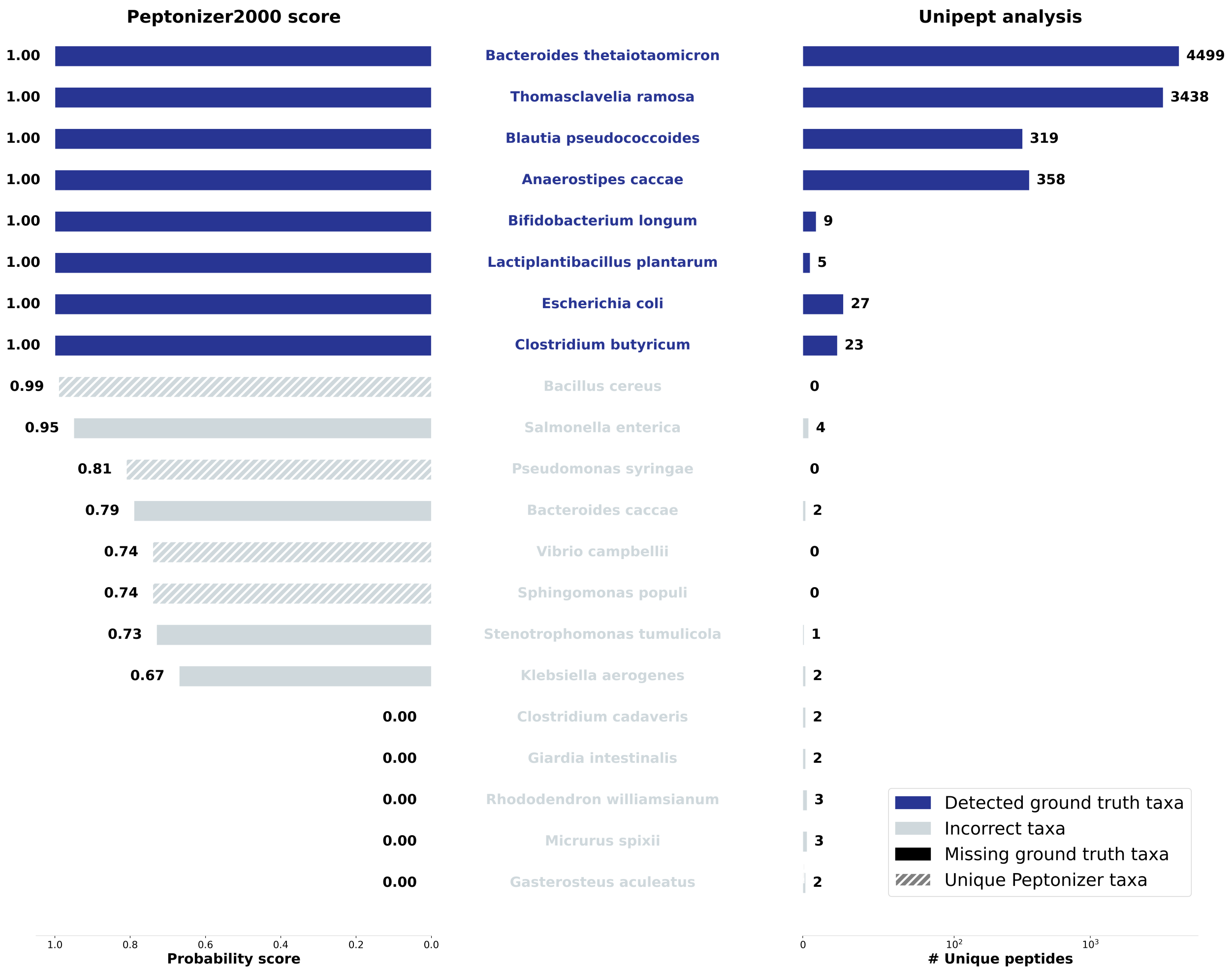


Figure 1: Results for the taxonomic identification of the SIHUMIx sample s3 using the Peptonizer2000, left, with corresponding probability scores, and using unique peptides from Unipept, right. Correctly identified taxa are shaded in a darker blue.

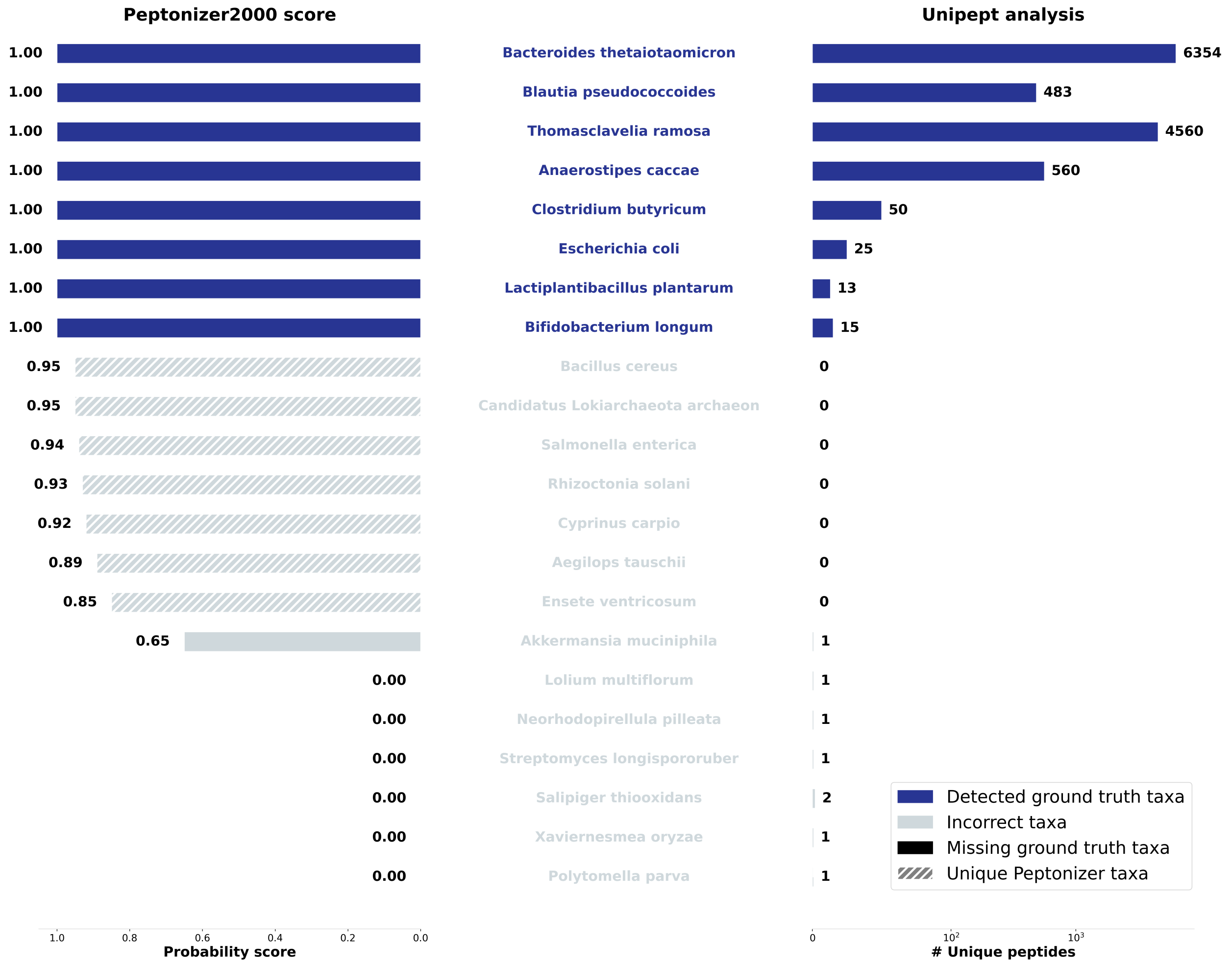


Figure 2: Results for the taxonomic identification of the SIHUMIx sample s5 using the Peptonizer2000, left, with corresponding probability scores, and using unique peptides from Unipept, right. Correctly identified taxa are shaded in a darker blue.

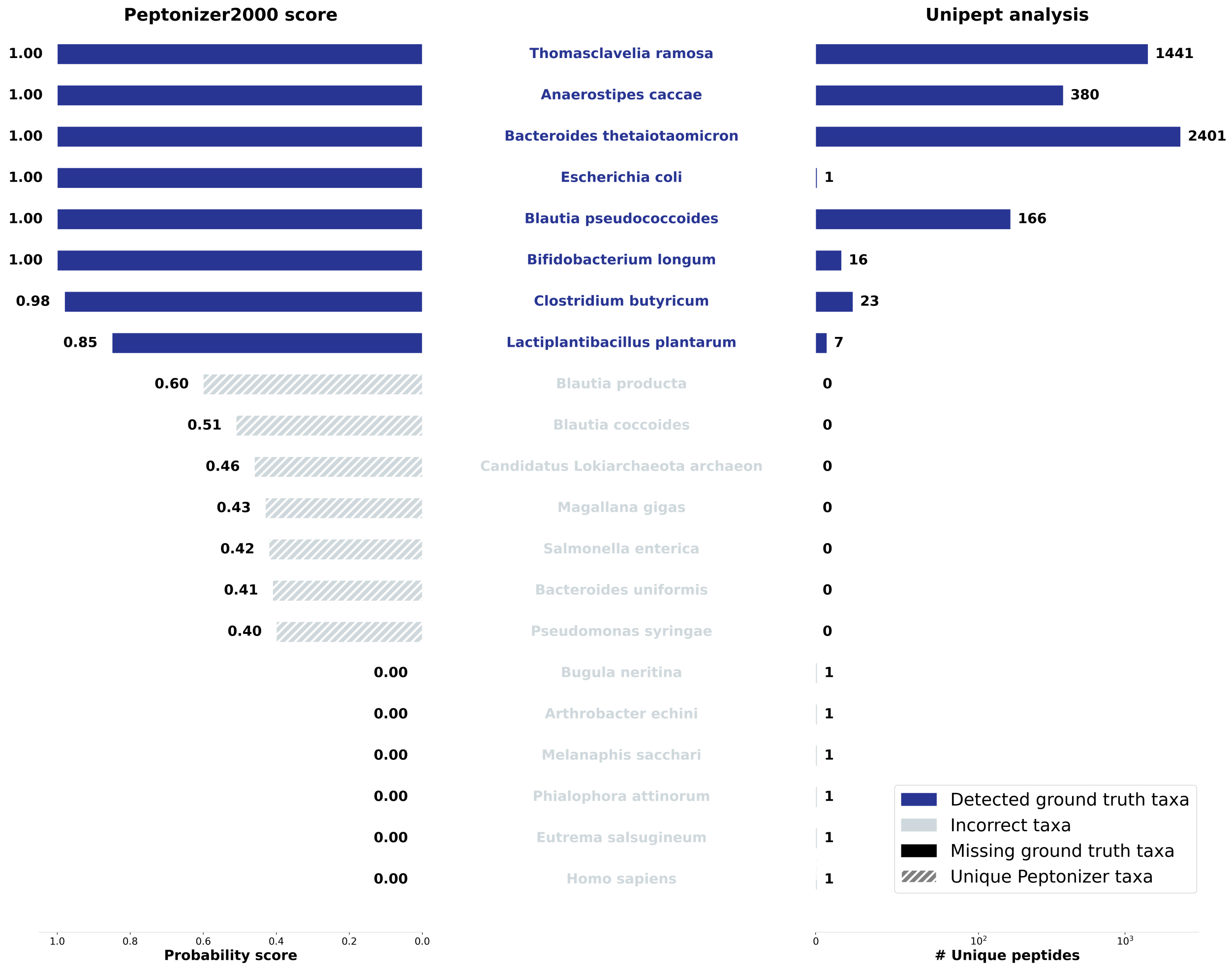


Figure 3: Results for the taxonomic identification of the SIHUMIx sample s8 using the Peptonizer2000, left, with corresponding probability scores, and using unique peptides from Unipept, right. Correctly identified taxa are shaded in a darker blue.

#### More complex samples

##### Lab assembled mixtures C1 and P1

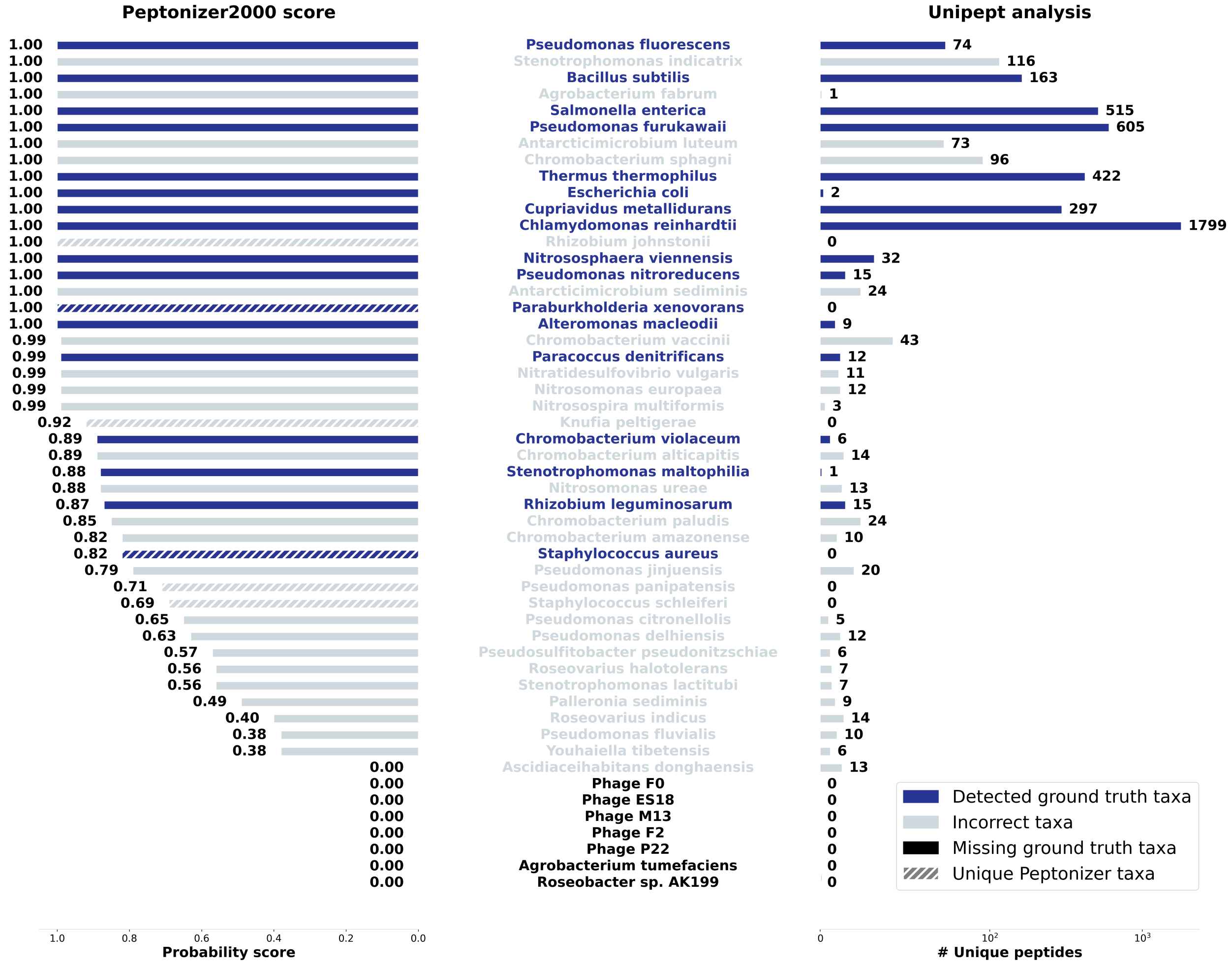


Figure 4: Results for the taxonomic identification of the lab assembled mixture with equal cell content C1 using the Peptonizer2000, left, with corresponding probability scores, and using unique peptides from Unipept, right. Correctly identified taxa are shaded in a darker blue.

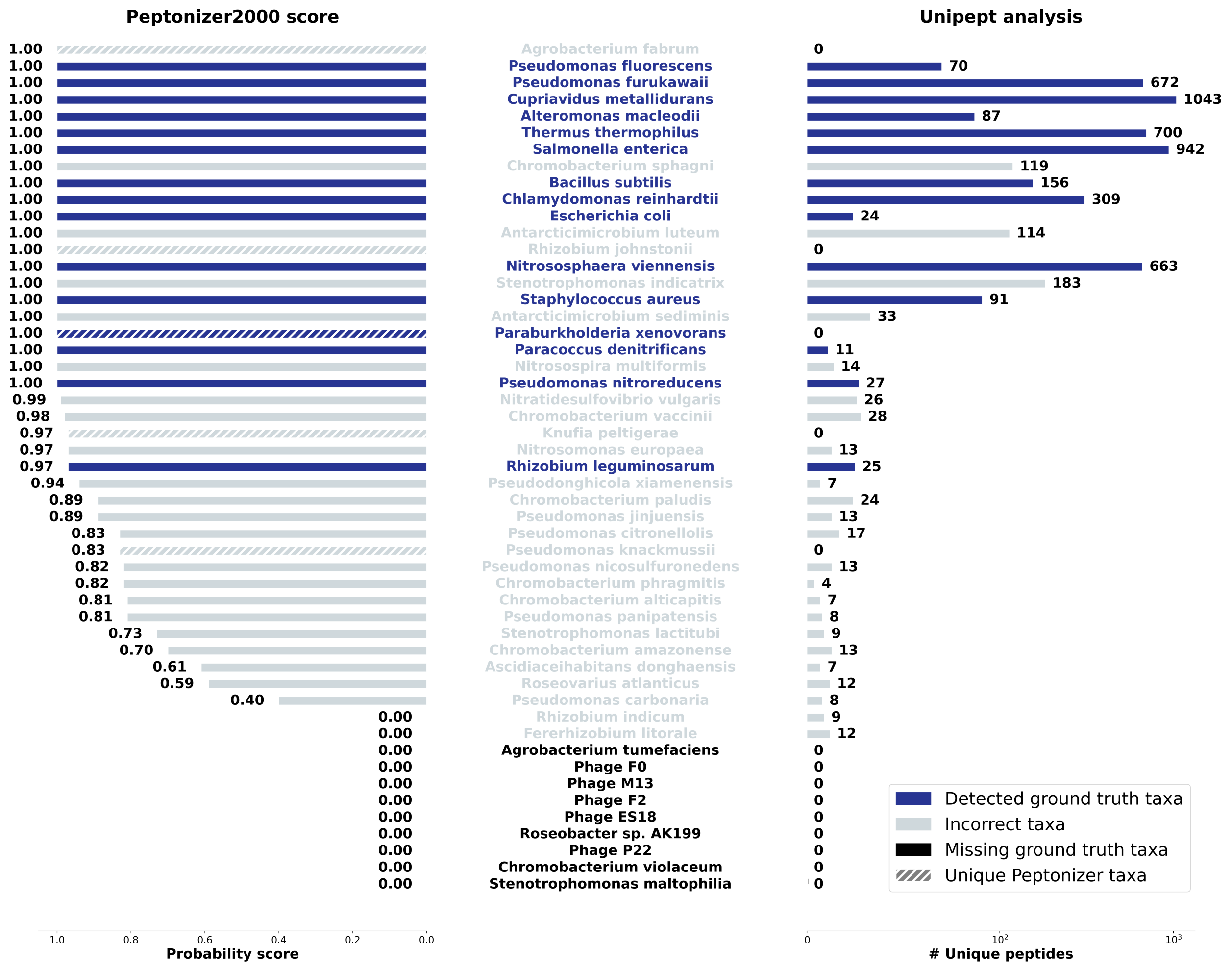


Figure 5: Results for the taxonomic identification of the lab assembled mixture with equal protein content P1 using the Peptonizer2000, left, with corresponding probability scores, and using unique peptides from Unipept, right. Correctly identified taxa are shaded in a darker blue.

#### Search against UNIREF50 and investigation of FDR effect

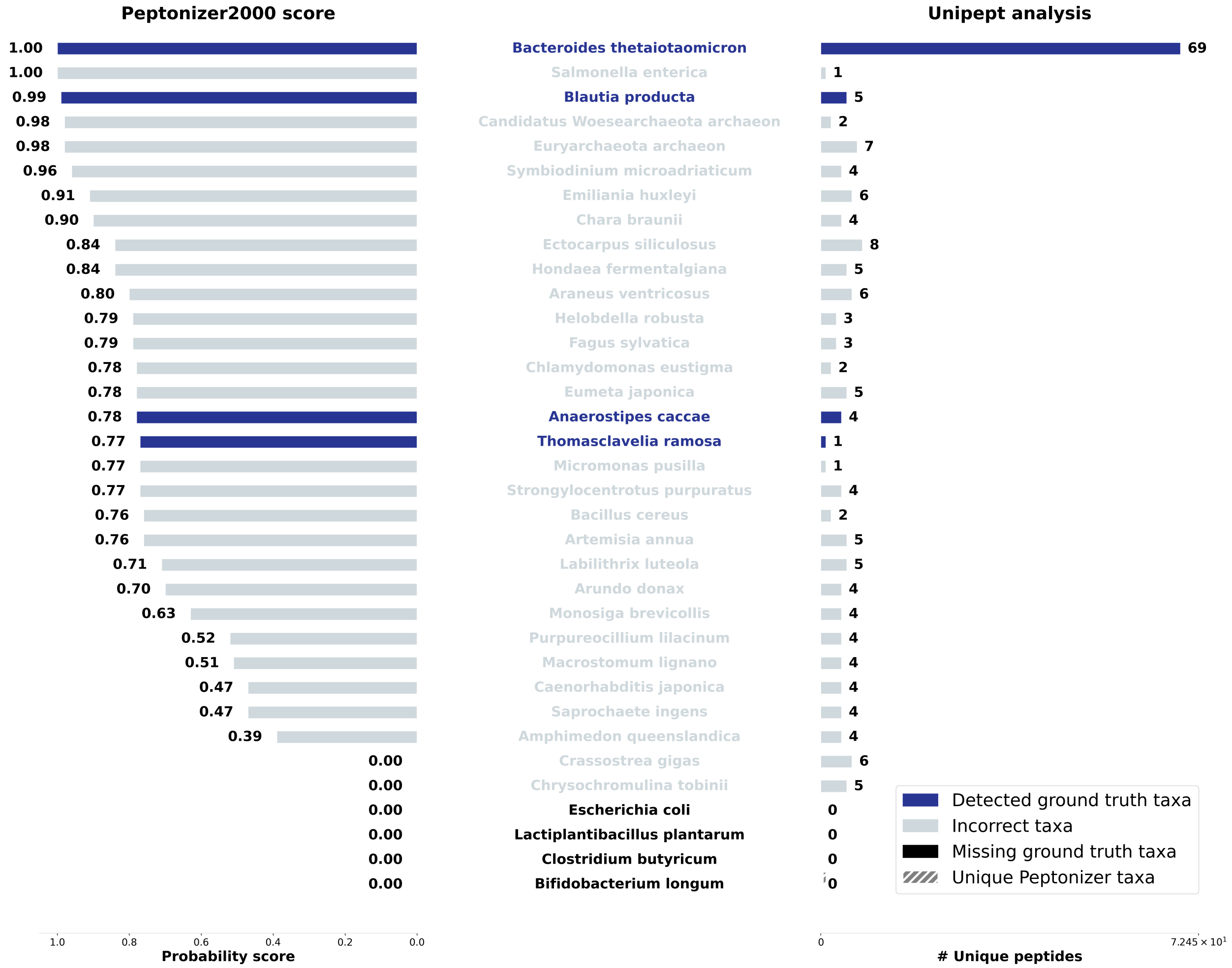


Figure 6: Results for the taxonomic identification of the SIHUMIx sample S07 searched against UNIREF50 with an FDR of 1%, using the Peptonizer2000, left, with corresponding probability scores, and using unique peptides from Unipept, right. Correctly identified taxa are shaded in a darker blue.

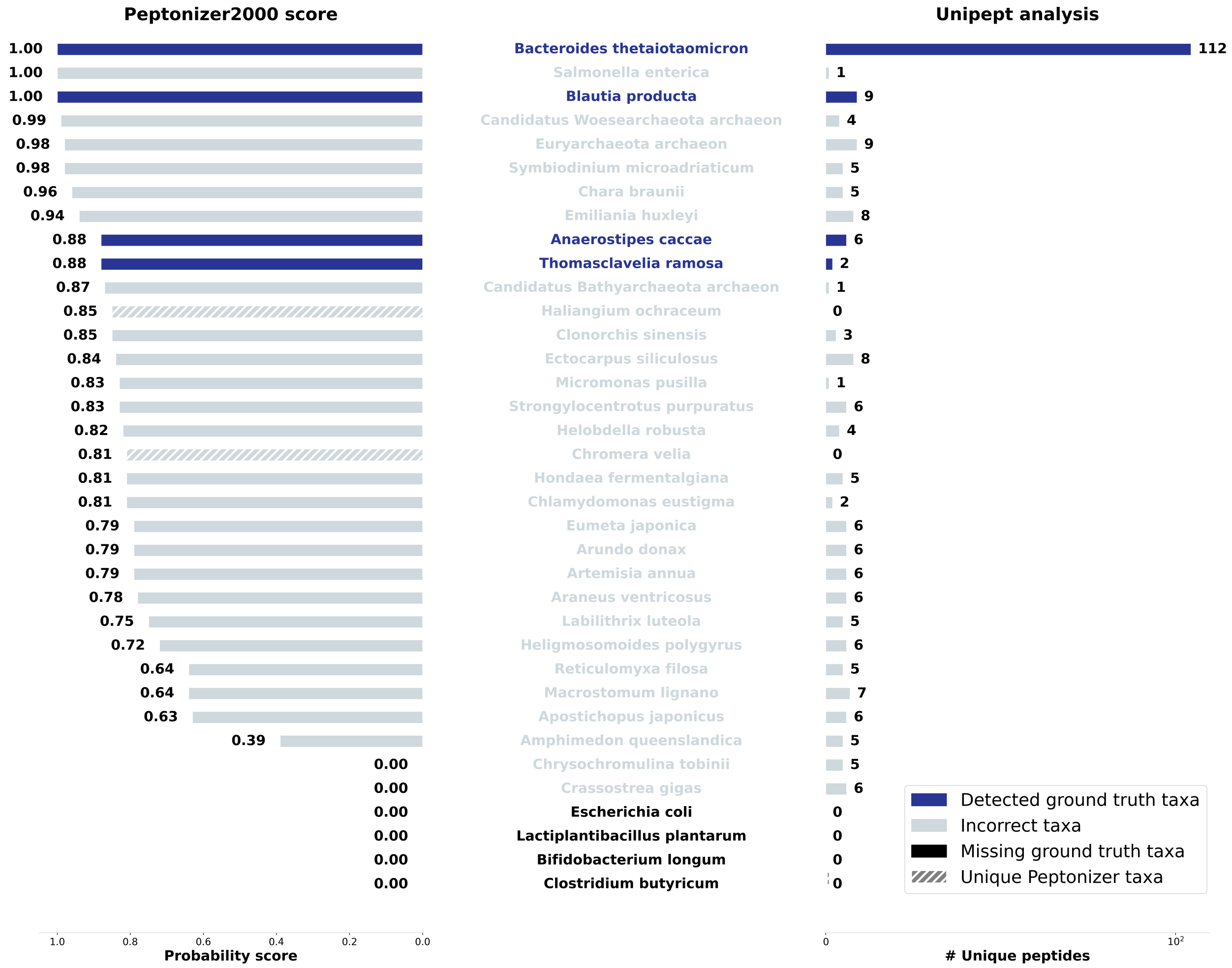


Figure 7: Results for the taxonomic identification of the SIHUMIx sample S07 searched against UNIREF50 with an FDR of 5%, using the Peptonizer2000, left, with corresponding probability scores, and using unique peptides from Unipept, right. Correctly identified taxa are shaded in a darker blue.

#### Fecal sample

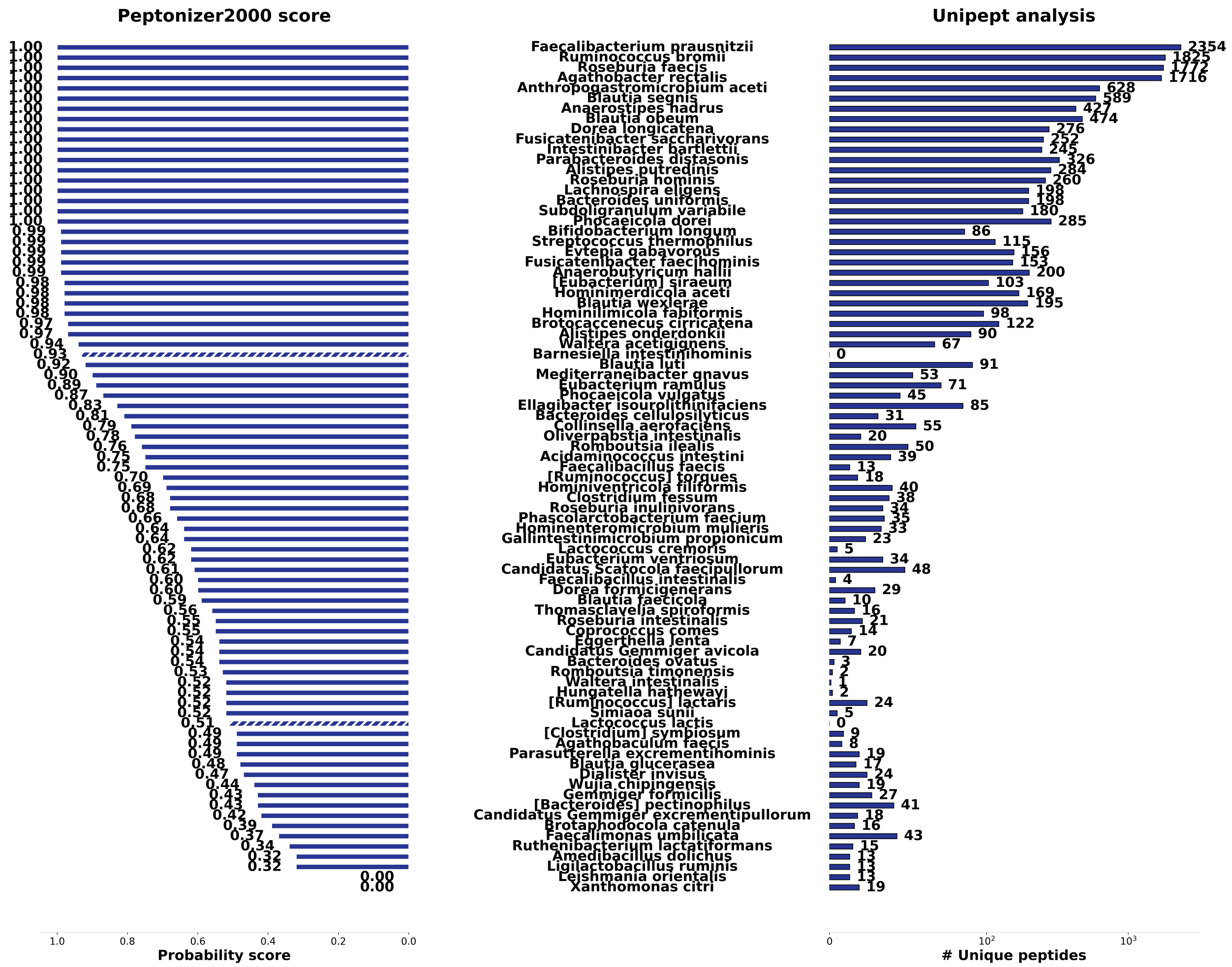


Figure 8: Results for the taxonomic identification of the fecal sample F07 at species level using the Peptonizer2000, left, with corresponding probability scores, and using unique peptides from Unipept, right.

### References


(1) Huerta-Cepas, J.; Serra, F.; Bork, P. ETE 3: Reconstruction, Analysis, and Visualization of Phylogenomic Data. *Molecular Biology and Evolution* **2016**, *33*, 1635–1638.

(2) McKinney, W.; others Data structures for statistical computing in python. Proceedings of the 9th Python in Science Conference. 2010; pp 51–56.

(3) Hagberg, A. A.; Schult, D. A.; Swart, P. J. Exploring Network Structure, Dynamics, and Function using NetworkX. Proceedings of the 7th Python in Science Conference. Pasadena, CA USA, 2008; pp 11 – 15.

(4) Virtanen, P. et al. scipy/scipy: SciPy 0.19.0. *Zenodo* **2017**, Publisher: Zenodo ADS Bibcode: 2017zndo....376244V.

(5) Cock, P. J. A.; Antao, T.; Chang, J. T.; Chapman, B. A.; Cox, C. J.; Dalke, A.; Friedberg, I.; Hamelryck, T.; Kauff, F.; Wilczynski, B.; de Hoon, M. J. L. Biopython: freely available Python tools for computational molecular biology and bioinformatics. *Bioinformatics* **2009**, *25*, 1422–1423.

(6) Hunter, J. D. Matplotlib: A 2D graphics environment. *Computing in Science & Engineering* **2007**, *9*, 90–95.

(7) Waskom, M. L. seaborn: statistical data visualization. *Journal of Open Source Software* **2021**, *6*, 3021.

(8) Kleiner, M.; Thorson, E.; Sharp, C. E.; Dong, X.; Liu, D.; Li, C.; Strous, M. Assessing species biomass contributions in microbial communities via metaproteomics. *Nature Communications* **2017**, *8*, 1558, Number: 1 Publisher: Nature Publishing Group.
